# Dual roles of mTORC1-dependent activation of the ubiquitin-proteasome system in muscle proteostasis

**DOI:** 10.1101/2021.11.16.468773

**Authors:** Marco S. Kaiser, Giulia Milan, Shuo Lin, Filippo Oliveri, Kathrin Chojnowska, Lionel A. Tintignac, Nitish Mittal, Christian E. Zimmerli, David J. Glass, Mihaela Zavolan, Daniel J. Ham, Markus A. Rüegg

**Affiliations:** Biozentrum, University of Basel, Basel, Switzerland; Novartis Institutes for Biomedical Research, Cambridge, MA, USA

**Keywords:** Nfe2l1, Nrf1, PKB/Akt, FoxO, proteasome, Atrophy, Ubiquitin-proteasome system

## Abstract

Muscle size is controlled by the PI3K-PKB/Akt-mTORC1-FoxO pathway, which integrates signals from growth factors, energy and amino acids to activate protein synthesis and inhibit protein breakdown. While mTORC1 activity is necessary for PKB/Akt-induced muscle hypertrophy, its constant activation alone induces muscle atrophy. Here we show that this paradox is based on mTORC1 activity promoting protein breakdown through the ubiquitin-proteasome system (UPS) by simultaneously inducing ubiquitin E3 ligase expression *via* feedback inhibition of PKB/Akt and proteasome biogenesis *via* Nuclear Factor Erythroid 2-Like 1 (Nrf1). Muscle growth was restored by reactivation of PKB/Akt, but not by Nrf1 knockdown, implicating ubiquitination as the limiting step. However, both PKB/Akt activation and proteasome depletion by Nrf1 knockdown led to an immediate disruption of proteome integrity with rapid accumulation of damaged material. These data highlight the physiological importance of mTORC1-mediated PKB/Akt inhibition and point to juxtaposed roles of the UPS in atrophy and proteome integrity.

## INTRODUCTION

The Protein Kinase B (PKB)/Akt-mTORC1-FoxO pathway is the central upstream signaling axis controlling skeletal muscle protein homeostasis (proteostasis), regulating both protein synthesis and breakdown, the latter *via* the autophagy/lysosomal system and the ubiquitin proteasome system (UPS). Activating PKB/Akt potently stimulates mTORC1-dependent protein synthesis and phosphorylates and sequesters FoxO transcription factors in the cytoplasm preventing their nuclear translocation and transcriptional activation of the UPS, thereby driving rapid skeletal muscle growth ^1^. However, there is feedback inhibition upon mTORC1 activation from S6K1 to IRS1 on PKB/Akt ^2^. Thus, constant activation of mTORC1 by genetic elimination of its upstream inhibitor TSC1, paradoxically not only fails to stimulate hypertrophy but induces atrophy in skeletal muscle, despite autophagy inhibition and the promotion of protein synthesis ^3, 4^. These observations implicate a parallel, mTORC1-mediated activation of protein breakdown pathways. Indeed, we have previously observed elevated levels of the key, atrophy-promoting E3 ubiquitin ligase *Trim63* (MuRF1) and the E3 ubiquitin ligase member *Fbxo32* (atrogin1/MAFbx) in muscle from mice with a muscle-specific knockout of the mTORC1 inhibitor *Tsc1* (TSCmKO mice) ^4^. The UPS is strongly activated during skeletal muscle atrophy and is considered the primary driver of muscle loss in many muscle-wasting conditions. On the other hand, it is the major system responsible for removing damaged proteins and has been shown to partially compensate for impaired autophagy and thereby to maintain cellular homeostasis ^5^. Since mTORC1 activation, autophagy inhibition and UPS induction are common features of numerous muscle-wasting conditions including denervation and sarcopenia ^4, 6^, understanding the underlying mechanisms responsible for mTORC1-mediated UPS induction and ultimately the consequence of its activity on muscle proteostasis is crucial to identifying strategies to counteract muscle wasting.

The UPS is a highly sophisticated proteolytic system that involves targeted labelling of substrates with ubiquitin *via* sequential transfer from the E1 ubiquitin activating enzyme to E2 ubiquitin-conjugating enzymes and then to one of more than 300 E3 ubiquitin ligases, which confer both substrate and ubiquitin chain conjugation specificity. For example, targets of the E3 ligase MuRF1 are thought to include major muscle structural proteins (e.g. titin, myosin heavy chains and some myosin light chains ^7^), while atrogin1/MAFbx is thought to target key proteins involved in protein synthesis (e.g. eIF3f ^8^).Once a substrate is labelled with a chain of four or more ubiquitin proteins (i.e. polyubiquitinated), it is sent to the functional unit responsible for UPS-induced protein degradation, which is referred to as the 26S proteasome based on its sedimentation coefficient. The 26S proteasome consists of the 20S catalytic core particle and one or two 19S regulatory particles. The 20S core particle is a cylindrical structure with a central pore made up of four stacked rings each containing seven individual proteasome subunit proteins. The inner two catalytic rings each contain PSMB1-7, while the outer two rings contain PSMA1-7 and act as a gate for substrate entry. The β5 (PSMB5), β1 (PSMB6), and β2 (PSMB7) catalytic enzymes are responsible for chymotrypsin-, caspase- and trypsin-like activity during protein cleavage, respectively. As the catalytic sites are located on the interior surface of the central pore, their activity relies on substrate access, a process mediated by 19S regulatory particles, which are responsible for recognizing and capturing (PSMD4, ADRM1), de-ubiquitinating (PSMD14) and unfolding ubiquitinated substrates before gate opening (PSMC1, 3 and 4) and substrate transport (PSMC1-6) via ATPase activity into the 20S core particle for destruction ^9^.

To better understand muscle proteostasis networks controlled by the PKB/Akt-mTORC1-FoxO pathways, we here investigated the mechanism underlying muscle wasting upon deletion of *Tsc1*. Based on transcriptomic and proteomic analysis, we find that muscles from TSCmKO mice increase expression of many atrophy-related genes (aka “atrogenes”) ^10, 11^, including many E3 ubiquitin ligases, as well as most subunits of the 26S proteasome, resulting in increased proteasome activity. Inducible deletion of *Tsc1* in adult skeletal muscle (iTSCmKO mice) for as little as 10 days was sufficient to reproduce atrogene expression, proteasome subunit expression and activity as well as muscle wasting in fast, but not slow-twitch muscles (e.g. *soleus*). By detailed examination of signaling pathways, genetic activation of PKB/Akt and knockdown experiments, we identified nuclear factor, erythroid derived 2-like 1 (Nfe2l1, also known as Nrf1) as the transcription factor responsible for 26S proteasome expression, while the PKB/Akt-FoxO axis was responsible for atrogene expression and muscle atrophy. Just 12 days of PKB/Akt activation induced a severe myopathy, including p62-positive aggregate accumulation and vacuoles in TSCmKO mice, despite potently inducing hypertrophy. Likewise, Nrf1-kd mediated proteasome depletion led to p62 accumulation and induction of atrogene expression. Together, these data point to juxtaposed roles of the UPS in skeletal muscle proteostasis, on the one hand driving muscle atrophy, but on the other alleviating cellular stress by removing misfolded and damaged proteins.

## RESULTS

### Sustained skeletal muscle mTORC1 activity upregulates the ubiquitin proteasome system

To understand the molecular mechanisms causing atrophy in muscles with high mTORC1 activation, we first analyzed the transcriptome of the *extensor digitorum longus* (EDL) muscle in 3-month-old TSCmKO mice ^6^. A short-term (3 day) treatment with rapamycin was used to identify transcripts that respond acutely to mTORC1 inhibition. Differential expression (DE) analysis identified 857 and 502 transcripts with significantly (FDR<0.05) higher and lower expression, respectively, in TSCmKO muscle compared to control that were also reversed by rapamycin (**Figure 1A, upper**). According to DAVID ^12^, the ‘biological process’ and ‘molecular function’ gene ontology (GO) terms that were most enriched in genes with higher expression in TSCmKO mice include multiple terms associated with the UPS (e.g. ‘protein ubiquitination’, ‘ubiquitin protein ligase activity’ and ‘ubiquitin-protein transferase activity’) as well as terms associated with ER stress and autophagy (**Figure 1A, lower**). GO terms enriched in genes that are repressed by mTORC1 were related to binding of RNA, calmodulin, metal ions, nucleic acids, ATP and nucleotides (**Figure 1A, lower**). Next, we compared the 857 upregulated ‘mTORC1-regulated’ genes with a curated list of 249 atrophy-related genes, known from literature to be increased during experimental paradigms (e.g., starvation, diabetes, cachexia, denervation) of muscle atrophy (**Table S1**). Remarkably, expression of 72 of the 249 atrophy-related genes were significantly (FDR<0.05) upregulated in TSCmKO mice and almost half of them (29) were normalized (FDR<0.05) by 3 days of rapamycin treatment (**Figure 1B**). Many of the genes in this class are involved in ubiquitin-proteasomal degradation, such as *Fbxo32* (atrogin1/MAFbx), *Trim63* (MuRF1), *Fbxo30* (MUSA), *Fbxo40, Mdm2, Traf6, Vcp (*p97*), Psmd8 and Psme4* (PA200) or in autophagy-lysosomal degradation, such as *Becn1, Bnip3, Ctsl, Fam134b, Gabarapl1* and *Sqstm1 (*p62) ^10, 13, 14, 15, 16, 17, 18, 19^.

**Figure 1:**
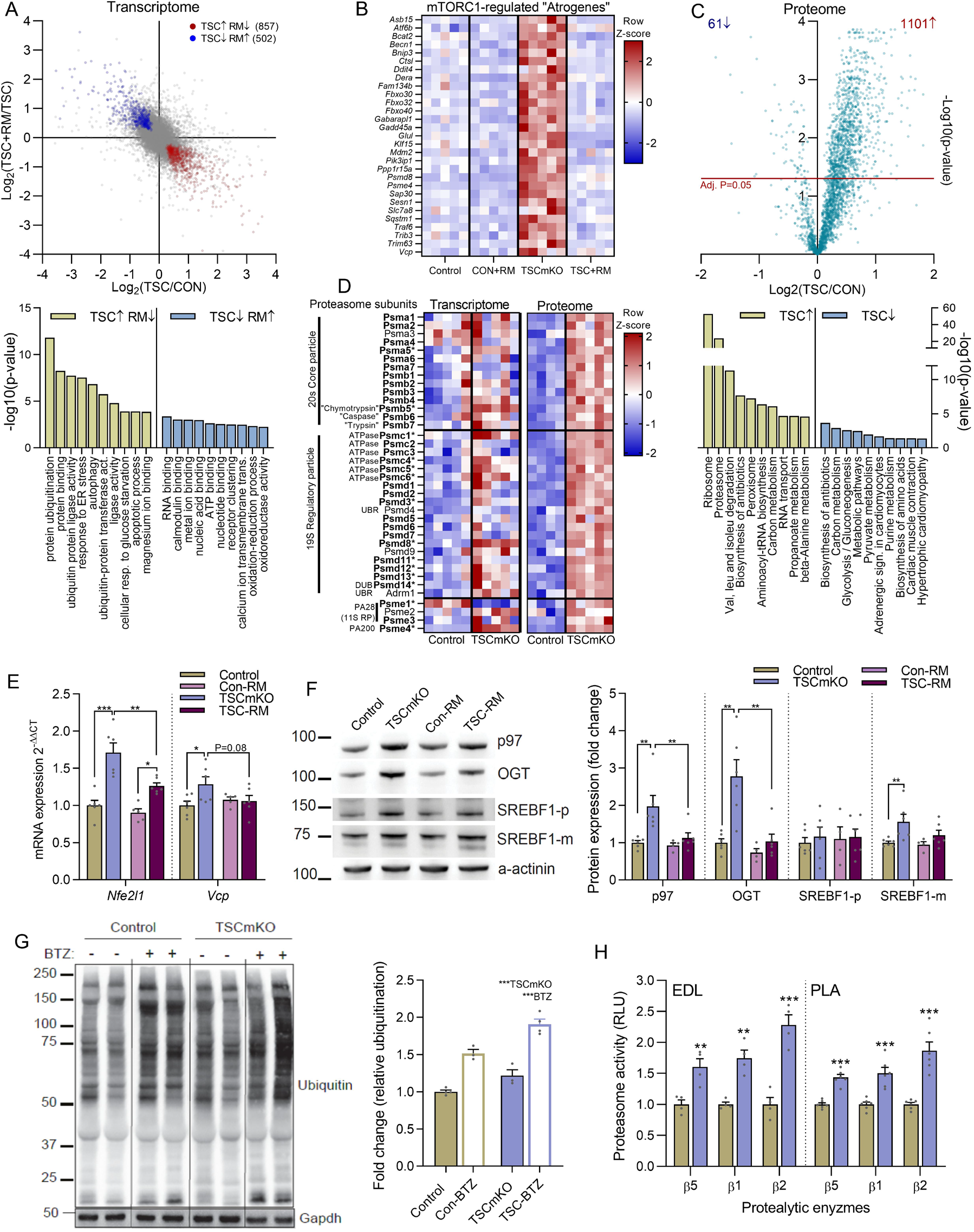
Skeletal muscle mTORC1 hyperactivity upregulates the ubiquitin proteasome system. **(A)** Pairwise comparison of muscle mTORC1 activation-(TSC/CON) and subsequent rapamycin inhibition-induced (TSC+RM/TSC) gene expression changes in EDL muscle with mTORC1-regulated genes up-(red) and downregulated (blue) in TSCmKO mice and counter-regulated by rapamycin as denoted. Top Gene Ontology (GO) terms are presented below (n=5 per group). **(B)** Heatmap of changes in the expression of mTORC1-regulated “atrogenes”. **(C)** Volcano plot for changes in protein expression between TSCmKO and control TA muscle with top GO terms associated with significantly increased and decreased proteins below (n=4-5 per group). **(D)** Heatmap of changes in gene and protein expression for control and TSCmKO mice of all expressed 26S proteasome subunits and members of the PA28 and PA200 proteasome activators. **(E)** mRNA and **(F)** protein expression of Nrf1 regulators, as measured by RT-qPCR (*gastrocnemius*) and Western blot (*tibialis anterior*), respectively (n=5-6 per group). **(G)** Representative blots and quantification of mono- and poly-ubiquitinated proteins in control and TSCmKO mice treated with vehicle or 1 mg.kg^-1^ of the proteasome inhibitor Bortezomib (BTZ) 12-16 h before dissection (n=3-4). **(H)** Luciferase-based peptidase activity of 20S proteasome catalytic enzymes in extensor digitorum longus (EDL; n=4) and plantaris (PLA; n=6) muscle. *Actb* was used as the reference gene (E) and either α-actinin (F) or GAPDH (G) was used as the protein loading control. Data are presented as mean ± SEM. Two-tailed Student’s t-tests (H) or two-way ANOVAs with Sidak post hoc tests were used to compare the data. *, **, and *** denote a significant difference between groups of *P* < 0.05, *P* < 0.01, and *P* < 0.001, respectively. For trends, where 0.05 < *P* < 0.10, p values are reported.

To evaluate whether the protein degradation signature revealed by transcript analysis was also observed at the protein level, we used mass spectrometry (MS) to identify differentially expressed proteins from *tibialis anterior* (TA) muscle of 3-month-old TSCmKO mice and littermate controls. DE analysis identified 1101 significantly (p<0.05) upregulated and 61 significantly downregulated proteins between control and TSCmKO mice (**Figure 1C, upper**). The ribosome and proteasome were overwhelmingly the most prominently enriched KEGG pathways (**Figure 1C, lower**). The ribosomal protein enrichment in TSCmKO mice is consistent with the prominent role of mTORC1 in the translation of ribosomal protein mRNA ^20, 21, 22^. Of the 14 and ∼21 proteins that make up the 20S core particle and the 19S regulatory particle of the 26S proteasome, 13 and 17 proteins, respectively were significantly increased in TSCmKO muscle compared to control (**Figure 1D, bold**), along with the proteasome activators Psme1, 3 and 4 (**Figure 1D**). Although less prominent, many 26S proteasome subunits and activators were also significantly upregulated (denoted with an asterisk) at the transcript level (**Figure 1D**). Confirming mRNA-seq data, RT-qPCR showed elevated expression of many atrogenes involved in ubiquitination, including the E3 ligases *Fbxo32, Trim63, Fbxo30, Mdm2* and *Traf6*, as well as the E3/E4 enzyme *Ube4b* and *Ubb* in TA muscle from 3-month-old TSCmKO mice compared to controls (**Figure S1A**). Since the UPS is the main protein degradation pathway activated during muscle atrophy ^23, 24^, we examined the regulation of this pathway in further depth. Confirming mRNA-seq and proteomics data, RT-qPCR (**Figure S1A**) and Western blot analysis (**Figure S1B**) showed elevated gene and/or protein expression of numerous proteasome subunits in TSCmKO mice, including catalytic (*Psmb5, Psmb6* and *Psmb7)* and non-catalytic (*Psma1* and *Psma5)* 20S core particle subunits as well as ATPase (*Psmc1*) and non-ATPase (*Psmd8* and *Psmd11*) 19S regulatory particle components along with the proteasome activator *Psme4*.

mTORC1 activation is high in old, sarcopenic muscle, independent of PKB/Akt, and rapamycin or rapalogs attenuate sarcopenia ^6, 25^, suggesting that aging may cause a similar gene expression signature as TSCmKO mice. To test this hypothesis, we analysed atrogene and proteasome subunit gene expression in previously generated transcriptomic data (https://sarcoatlas.scicore.unibas.ch/) from *tibialis anterior* muscle of adult (10mCON), 30-month-old, sarcopenic (30mCON), and 30-month-old, rapamycin-treated mice (30mRM) ^6^. Although changes were less strong in the aging data set, we saw a remarkably similar response to that seen in TSCmKO mice with approximately two thirds of mTORC1-regulated atrogenes (**Figure S2A**) and over 80% of 26S proteasome subunits (**Figure S2B**) being either higher (or tending to be higher) in 30mCON than 10mCON or lower (or tending to be lower) in 30mRM than 30mCON, or both. These data thus indicate that sustained activation of mTORC1 at high age exerts similar responses as its genetic activation by depletion of TSC1.

The remarkably consistent and ubiquitous upregulation of proteasome subunits at both the transcript and protein level suggests the involvement of a coordinating transcription factor. While FoxO transcription factors are known to control expression of many of the mTORC1-regulated ubiquitin- and autophagy-related atrophy genes, they are not known to broadly regulate the expression of proteasome subunits. Although currently uncharacterized in skeletal muscle, the nuclear factor E2-related factor 1 (Nfe2l1 or Nrf1) transcription factor has been identified as a master regulator of proteasome subunit gene expression, binding to the core antioxidant response element (ARE) sequence in the promoter region of all proteasome subunit genes ^26, 27, 28, 29, 30, 31^. Consistent with its involvement in the transcriptional regulation of the UPS in TSCmKO mice, mRNA expression of *Nfe2l1* and *Vcp*, which encodes the protein p97 and is responsible for Nrf1 transport from the ER lumen into the cytosol ^17, 32^, were significantly increased in TSCmKO muscles and reversed by 3 days of rapamycin treatment (**Figure 1E**). Similarly, protein levels of the Nrf1 regulators p97, O-linked N-acetyl glucosamine (O-GlcNAc) transferase (OGT), which mediates O-GlcNAcylation and stabilization of Nrf1 ^33, 34^, and the mature form of SREBF1, which mediates mTORC1-dependent induction of Nrf1 ^35, 36^, were all significantly higher in TSCmKO than control mice. Three days of rapamycin treatment normalized protein levels of p97, OGT and the mature form of SREBF1 in TSCmKO mice, indicating that mTORC1 activity is responsible for the increase in Nrf1 regulators (**Figure 1F**). In line with our observations, studies in cultured cells, brain and liver tissue have shown that genetic or physiological activation of mTORC1 increases *Nfe2l1* transcript levels and proteasome content through SREBF1 ^36^.

Despite the increase in proteasome content, ubiquitinated proteins were still higher in protein lysates from TSCmKO than control mice (**Figure 1G**). Ubiquitinated protein accumulation was not the result of impaired proteasome degradation as both control and TSCmKO mice treated with the proteasome inhibitor Bortezomib displayed a similar increase in ubiquitinated proteins (**Figure 1G**). Finally, a significant increase in 20S peptidase activity for all three proteolytic enzymes (i.e, chymotrypsin-like/β5, caspase-like/β1 and trypsin-like/β2) was seen in both EDL and plantaris (PLA) muscles of TSCmKO mice compared to controls (**Figure 1H**). Altogether, these data show that sustained mTORC1 activity promotes an atrophy-like transcriptional program in skeletal muscle ^11^, inducing the expression of atrophy-related E3 ligases as well as Nrf1 and a plethora of its 26S proteasome subunit targets, leading to increased proteasome activity.

### Acute TSC1 depletion rapidly activates the UPS in adult fast-twitch muscle

To further confirm that UPS induction in TSCmKO mice is a direct consequence of mTORC1 activation and not muscle adaptations to prolonged *Tsc1* deletion, we next examined mice in which recombination of the floxed *Tsc1* allele in skeletal muscle could be triggered by tamoxifen injection ^37, 38, 39^. To assure successful recombination, mice also carried an EGFP-reporter only expressed after Cre recombinase-mediated removal of a stop cassette ^37, 40^. We analyzed control and inducible TSCmKO (iTSCmKO) mice 10 (10d-iTSCmKO) or 21 (21d-iTSCmKO) days after tamoxifen-induced recombination (**Figure S3A**). 10d and 21d-iTSCmKO experiments were performed separately, normalized to their respective controls and then, based on comparable values and within-group variation, control groups from both experiments were pooled for ease of visualization and statistical comparisons. Ten days after recombination, all gastrocnemius muscle fibers were brightly GFP positive, indicating successful recombination (**Figure S3B**). In 10d-iTSCmKO muscle, TSC1 protein levels were significantly reduced and the mTORC1 targets S6 (S240/S244 and S235/S236) and 4E-BP1 (S65) were strongly phosphorylated, while the inhibitory feedback loop from S6K to PKB/Akt was not yet fully established with lower PKB/Akt (S473) but not PRAS40 (T246) phosphorylation (**Figure 2A**). After 21 days, phosphorylation of mTORC1 targets S6 and 4EBP1 was fully established, as was the dampening of PKB/Akt and PRAS40 phosphorylation (**Figure 2A**). The activation of mTORC1 and inhibition of PKB/Akt and PRAS40 correlated well with progressive fast-twitch muscle mass loss and slow-twitch muscle mass gain, as previously observed ^3, 4^, reaching statistical significance by 10 days in the TA and by 21 days in the *soleus* (SOL; **Figure 2B**).

**Figure 2:**
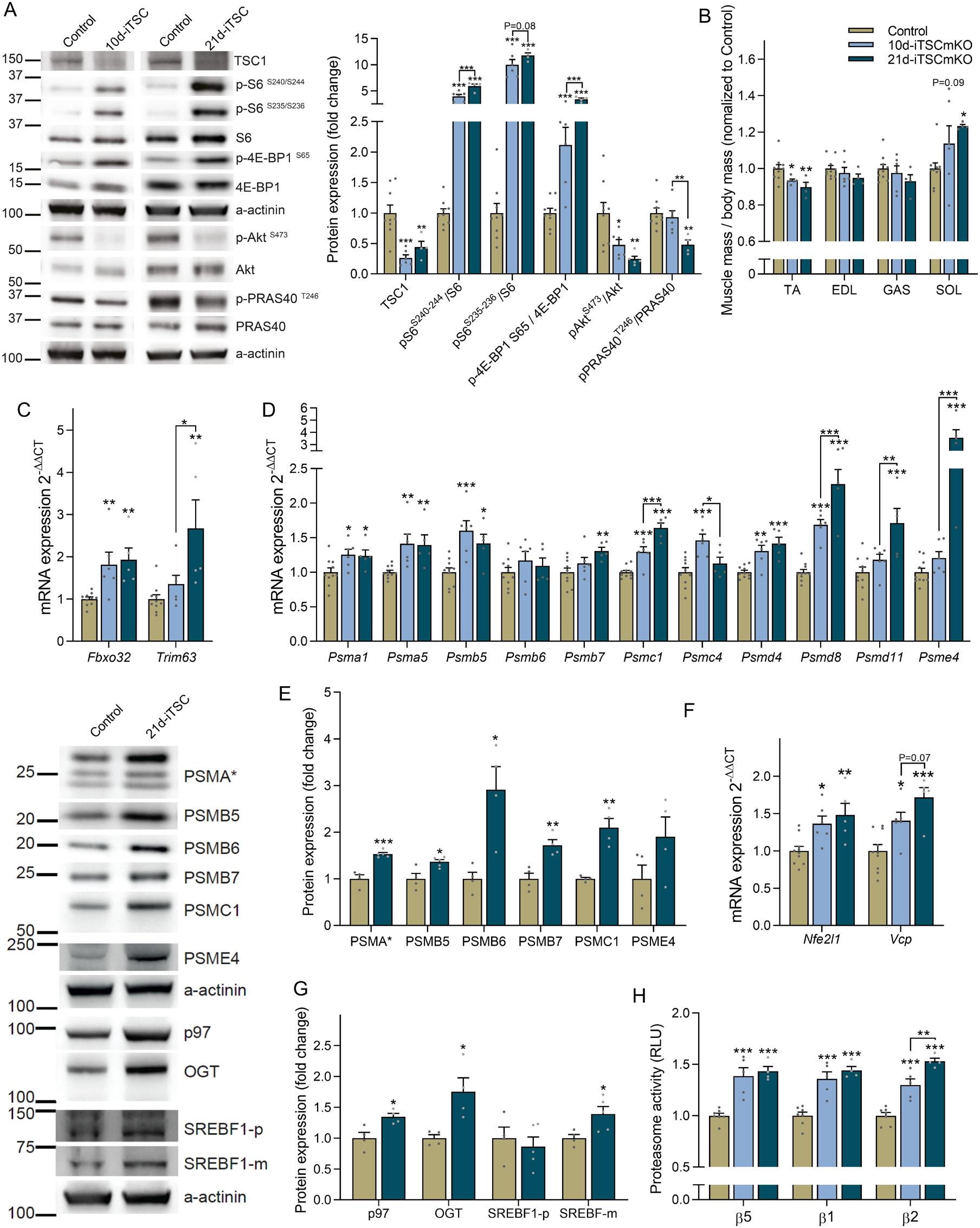
Acute TSC1 depletion rapidly activates the UPS in adult fast-twitch muscle. **(A**) Representative Western blots and quantification of proteins involved in mTORC1 signaling (*tibialis anterior*) and **(B)** muscle mass after 10 and 21 days recombination of floxed *Tsc1* alleles. Muscle mass of *tibialis anterior* (TA), *extensor digitorum longus* (EDL), *gastrocnemius* (GAS) and *soleus* (SOL) was averaged across both limbs, normalized to body mass and then to control. RT-qPCR measures of mRNA expression for **(C)** E3 ligases and **(D)** selected proteasome subunits after 10 and 21 days of *Tsc1* deletion (*gastrocnemius*). **(E)** Representative Western blots (left, upper) and quantification of 26S proteasome subunits and its activator (*tibialis anterior*). **(F)** mRNA and **(G)** protein expression (representative blots left, lower) for Nrf1 and its regulators, as measured via RT-qPCR and Western blot, respectively, after 10 and/or 21d of *Tsc1* deletion. **(H)** Luciferase-based peptidase activity of 20S proteasome catalytic enzymes. RT-qPCRs were performed on *gastrocnemius*, Western blots on *tibialis anterior* and proteasome activity on *extensor digitorum longus. Tubb* (C, D) or *Actb* (F) were used as reference genes, while α-actinin was used as a protein loading control. For A-D, F and H n=9 (Control), 6 (10d-TSCmKO) and 5 (21d-TSCmKO). For E and G, n=4-5. Data are presented as mean ± SEM. Two-tailed Student’s t-tests (E and G) or one-way ANOVAs (A-D, F and H) with Fishers LSD post hoc tests were used to compare the data. *, **, and *** denote a significant difference between groups of *P* < 0.05, *P* < 0.01, and *P* < 0.001, respectively. For trends, where 0.05 < *P* < 0.10, p values are reported.

Supporting a rapid mTORC1-mediated UPS induction, *Fbxo32* and *Trim63* were upregulated in both 10d-and 21d-iTSCmKO muscle (**Figure 2C**), along with a coordinated upregulation of 26S subunit genes after 10 days and a further increase after 21 days for some 19S regulatory particle subunits (*Psmc1, Psmd8* and *Psmd11*) as well as the proteasome activator *Psme4* (**Figure 2D**). We also confirmed the upregulation of protein expression for key 26S proteasome subunits in 21d-iTSCmKO mice (**Figure 2E**). In support of the mTORC1-induced upregulation of the proteasome being mediated by Nrf1, upregulation of *Nfe2l1* and *Vcp* (**Figure 2F**), along with the protein expression of the Nrf1 regulators p97, OGT and mature SREBF1 (**Figure 2G**), mirrored proteasome subunit expression in 10d- and/or 21d-iTSCmKO muscle. As in TSCmKO muscle, peptidase activity of the three catalytic β-proteasome subunits was strongly increased in muscle of 10d- and further increased in 21d-iTSCmKO mice compared to controls (**Figure 2H**).

As the slow-twitch muscle increases its size in response to sustained mTORC1 activity ^3^ and this increase in mass is already observed after short-term mTORC1 activation (**Figure 2B**), we also examined signaling in SOL muscle of 21d-iTSCmKO mice. The direct targets of mTORC1 as well as PBK/Akt and PRAS40 were affected as in fast-twitch muscles (**Figure S4A**). However, mRNA levels of two FoxO targets *Fbxo32* and *Trim63* (**Figure S4B**) as well as *Nfe2l1* and *Vcp* were not different between SOL muscles from 21d-TSCmKO and control mice (**Figure S4C**). Similarly, expression of mRNA (**Figure S4D**) and protein of the majority of the 26S proteasome subunits (**Figure S4E**) was substantially blunted or did not differ in SOL muscles between 21d-iTSCmKO and control mice. Importantly, there was also no difference in the proteolytic activity of the proteasomes (**Figure S4F**). These results show that the regulation of UPS components in response to mTORC1 activation differs between fast- and slow-twitch muscles and provides a possible explanation for the differential muscle mass response.

Together, these data demonstrate that mTORC1 rapidly activates the UPS, including atrophy-related ubiquitin E3 ligases and coordinate upregulation of the 26S proteasome and its transcriptional regulator Nrf1 in fast-twitch muscles as little as 10 days after floxed *Tsc1* allele recombination, which correlates with muscle atrophy. Hence, UPS activation is likely a direct result of mTORC1 activation rather than a consequence of the myopathic features associated with prolonged *Tsc1* deletion ^4^. UPS activation may also explain why fast-twitch muscles from TSCmKO mice are atrophic. However, these data cannot distinguish between the possibility that this phenotype is based on a PKB/Akt-FoxO-regulated increase in atrophy-related gene expression or an Nrf1-induced upregulation of 26S proteasome content, or both. To answer this, we next examined the effect of proteasome activity on skeletal muscle.

### Proteasome inhibition fails to rescue mTORC1-mediated muscle atrophy

As a first step, we directly targeted proteasome activity. We reasoned that if the mTORC1-induced increase in proteasome activity overwhelms the mTORC1-induced increase in protein synthesis to drive muscle wasting then perturbing proteasome activity should shift proteostatic balance and increase muscle mass in TSCmKO mice. To test this, we directly impaired proteasome function with the chemical proteasome inhibitor Bortezomib, which has been shown to target the β5 and to a lesser extent the β1 catalytic enzymes of the proteasome ^41^.

Based on previous experiments showing beneficial effects of systemic Bortezomib administration in mouse models of muscular dystrophy ^42^, we administered 1 mg/kg Bortezomib to control and TSCmKO mice every 72 h for four weeks and collected tissue 6 h following the final injection. Bortezomib-treated control and TSCmKO mice tended to gain less mass than vehicle-treated mice across the 4-week treatment period (**Figure 3A**), with adipose tissue impairments appearing to be the primary contributor to lower body mass increases in Bortezomib-treated control and TSCmKO mice (**Figure 3B**). Endpoint analysis showed that Bortezomib lowered muscle mass independent of genotype (**Figure 3C**). Fiber type-specific analysis of muscle fiber size showed a significant reduction in type IIX median fiber size in control mice and an enlargement of type IIA fibers in TSCmKO mice (**Figure 3D**), reminiscent of the hypertrophy seen in the slow-twitch soleus muscles of TSCmKO mice. Type IIA fiber enlargement has also been associated with the myopathic responses to denervation seen in TSCmKO mice ^39^. In line with this interpretation, Bortezomib failed to rescue the strong reduction in body-mass-normalized *ex vivo* peak tetanic EDL muscle force in TSCmKO mice (**Figure 3E**).

**Figure 3:**
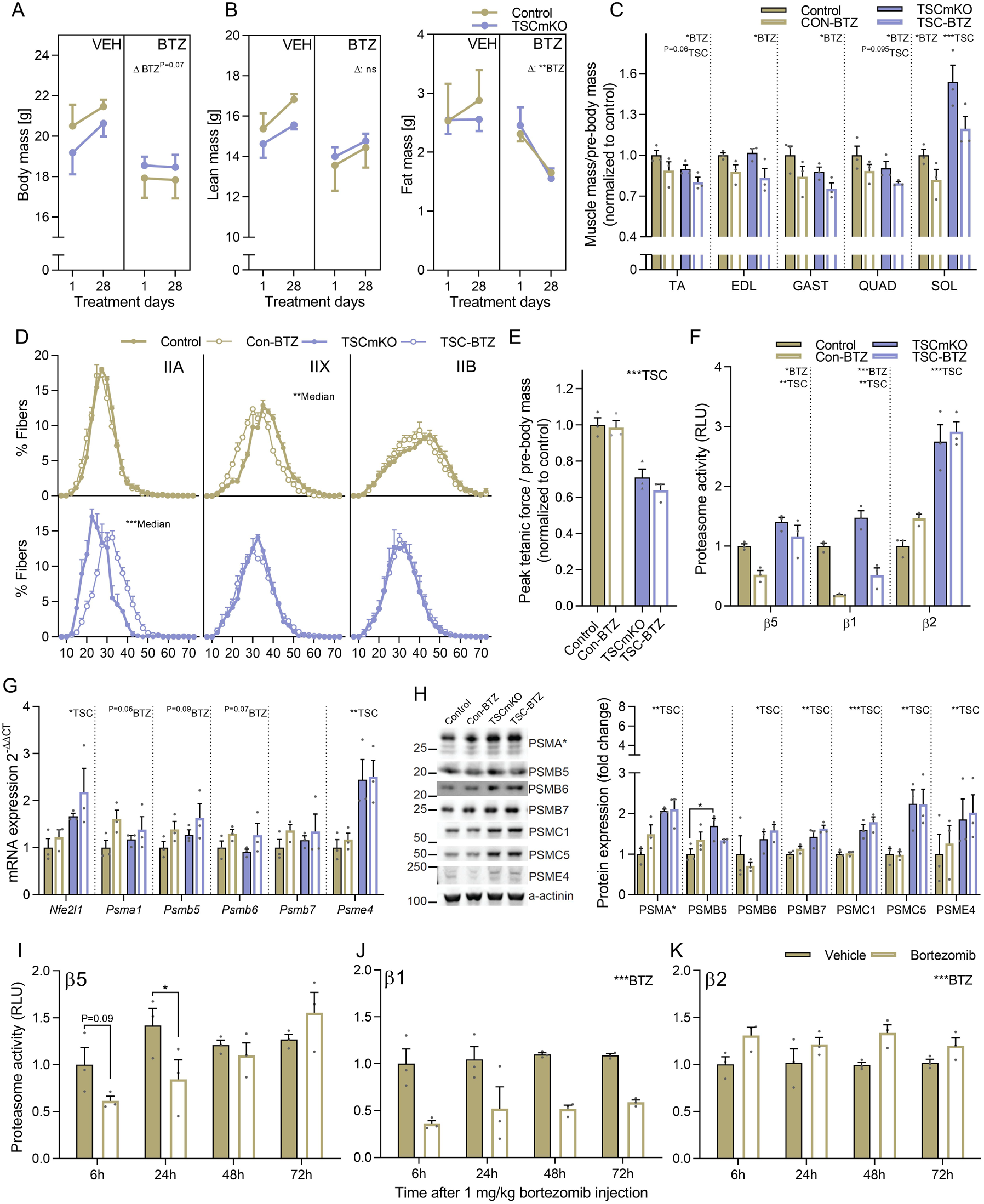
Proteasome inhibition via Bortezomib does not prevent mTORC1-induced muscle atrophy. Pre and post measures of **(A)** body, **(B)** lean and fat mass for vehicle and Bortezomib (BTZ; 1 mg.kg^-1^ every 72h) treated control and TSCmKO mice. **(C)** Muscle mass normalized to body mass **(D)** fiber-type specific minimum fiber feret distribution and **(E)** body mass normalized peak *ex vivo* tetanic force in *extensor digitorum longus* after 28d of vehicle or Bortezomib (BTZ) treatment in control (Con) and TSCmKO (TSC) mice. **(F)** Luciferase-based peptidase activity of 20S proteasome catalytic enzymes in *extensor digitorum longus* muscle after 28d Bortezomib treatment and 6 h after the final Bortezomib injection. **(G)** mRNA and **(H)** protein expression for proteasome subunits (and *Nfe2l1* mRNA) as measured by RT-qPCR and Western blot, respectively, after 28d of vehicle or Bortezomib treatment in control and TSCmKO mice. Luciferase-based peptidase activity of the 20S proteasome catalytic enzymes **(I)** β5, **(I)** β1 and **(J)** β2 in *extensor digitorum longus* (EDL) muscle 6, 24, 48 and 72h after a single injection of 1 mg.kg^-1^ Bortezomib. *Actb* was used as the reference gene and α-actinin as the protein loading control. For all experiments, n=3 per group. Data are presented as mean ± SEM. Two-tailed Student’s t-tests (D and I-K) or two-way ANOVAs with Sidak post hoc tests (A-C and E-H) were used to compare the data. *, **, and *** denote a significant difference between groups of *P* < 0.05, *P* < 0.01, and *P* < 0.001, respectively. For trends, where 0.05 < *P* < 0.10, p values are reported.

To confirm that Bortezomib impaired proteasome function in muscle after prolonged treatment, and in TSCmKO mice, we measured β5, β1 and β2 catalytic enzyme activity in EDL muscle. Bortezomib strongly suppressed β1 activity, slightly suppressed β5 activity, but did not affect β2 activity, independent of genotype (**Figure 3F**). Next, we compared transcript levels of Nrf1 and 20S proteasome subunits in the different samples. In all the Bortezomib-treated samples, *Nfe2l1* expression along with mRNA (**Figure 3G**) but not protein levels (**Figure 3H**) of the proteasome subunits tended to be increased, which is consistent with the well-documented Nrf1-mediated ‘bounce-back response’ of proteasome subunits upon proteasome inhibition ^43^. Since Bortezomib was administered every 72 hours in the 4-week-long treatment and endpoint analysis was performed 6 hours after the last injection, we next wanted to determine the extent to which proteasome activity inhibition was maintained over this time frame in mouse muscle. To this end, muscle tissue from control mice was collected 6, 24, 48 and 72 after a single Bortezomib injection. Similar to prolonged treatment, Bortezomib (1 mg/kg) reduced β5 chymotrypsin-like activity 6 (trend) and 24 h (P<0.05) after administration, but returned to control levels 48 and 72 h after the injection (**Figure 3I**). On the other hand, stronger and more prolonged inhibition of β1 caspase-like activity (**Figure 3J**) and a compensatory increase in β2 activity (**Figure 3C**), which Bortezomib is known not to effectively target ^41^, was observed across the full 72 h period. While these data show that Bortezomib administration successfully enters skeletal muscle tissue and impairs proteasome function with the effects more pronounced on β1 than β5 catalytic activity, they also highlight that more frequent administration would be necessary to maintain β5 activity inhibition. Since our attempts to administer Bortezomib more frequently compromised mouse survival, we decided that a more direct and muscle-specific approach to proteasome inhibition was required to conclusively test the role of higher proteasome content and activity in mTORC1-mediated atrophy.

### Nrf1 mediates the mTORC1-induced upregulation of the proteasome

To directly test how proteasome content affects muscle size, we next targeted *Nfe2l1* with small hairpin RNA (shRNA). As muscle contains many non-muscle fiber cells that could also contribute to the increased expression of *Nfe2l1* in whole-muscle lysates, we first used single molecule fluorescent *in situ* hybridization (smFISH; RNAscope®) to localize *Nfe2l1* expression. Indeed, the majority of *Nfe2l1* transcripts were expressed within skeletal muscle fibers in control mice and TSCmKO mice showed a strong increase of *Nfe2l1* puncta in the cytoplasm of the muscle fiber with sporadic, peripherally-localized clusters (**Figure 4A**). These data show that muscle fiber mTORC1 activation triggers the increase in *Nfe2l1* mRNA within muscle fibers and supports the notion that *Nfe2l1* expression is a direct consequence of mTORC1 activation.

**Figure 4:**
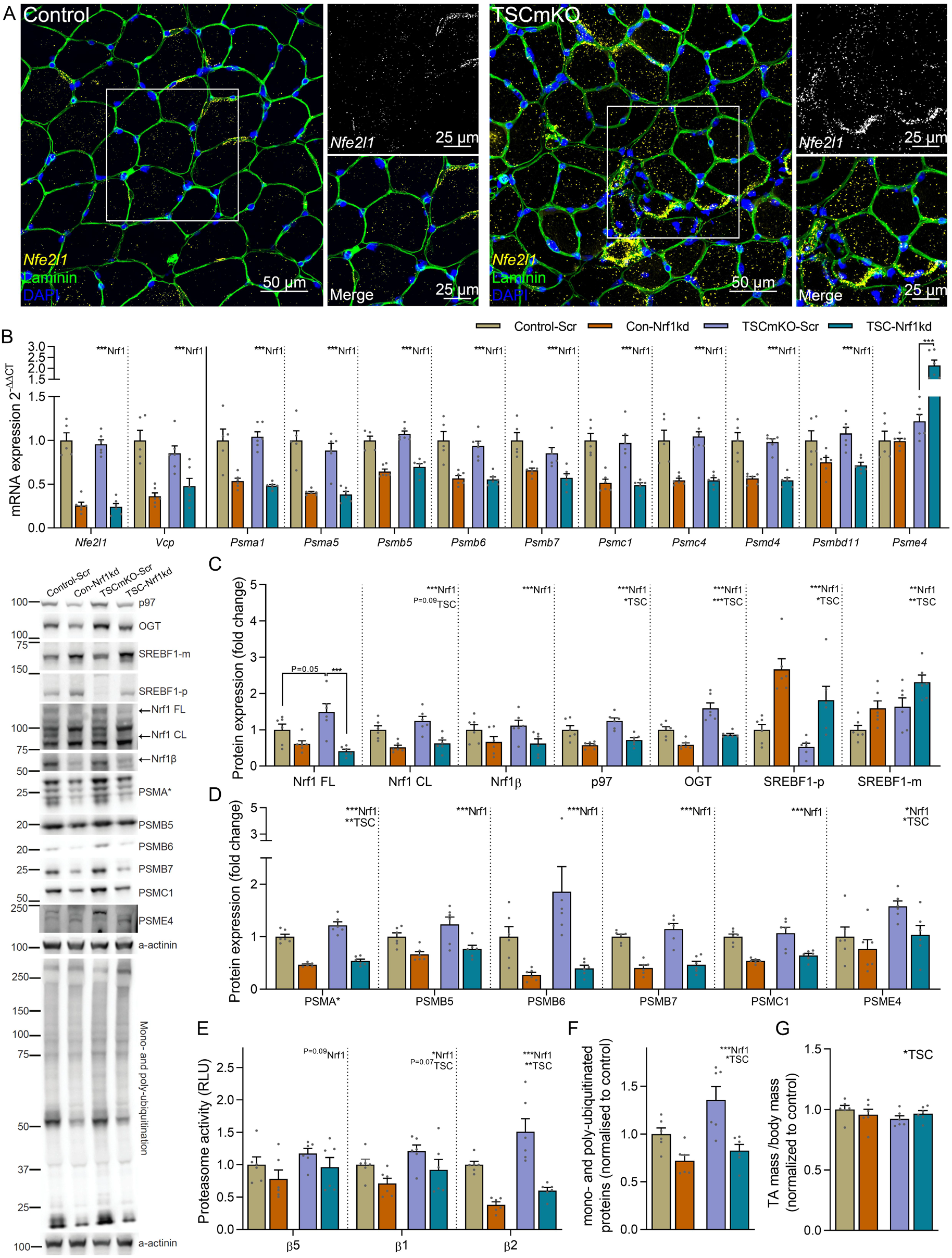
Nrf1 mediates mTORC1-stimulated proteasome biogenesis. **(A)** localization of *Nfe2l1* mRNA within muscle fibers using RNA *in situ* hybridization coupled to antibody-based immunofluorescence. **(B)** mRNA and protein expression **(C)** of Nrf1, encoded by the *Nfe2l1* gene, and Nrf1 regulators and **(D)** 26S proteasome subunits and the 20S activator PSME4 in *tibialis anterior* (TA) muscle with representative Western blots (left). Nrf1 FL, CL and β refer to the full length, cleaved and alternatively spliced versions of Nrf1, respectively. Muscles were collected 2 weeks after transfection of a plasmid expressing either a scrambled (Scr) or *Nfe2l1*-targeting (Nrf1kd) shRNA into TA/*extensor digitorum longus* (EDL) muscles of control and TSCmKO mice. **(E)** Luciferase-based peptidase activity of 20S proteasome catalytic enzymes in EDL muscle, **(E)** mono- and poly-ubiquitinated proteins and representative Western blots (left) and **(G)** TA muscle mass 2 weeks after transfection. *Des* was used as the reference gene and α-actinin as the protein loading control. For all experiment n=6 per group, except for control in B where n=5. Data are presented as mean ± SEM. Two-way ANOVAs with Sidak post hoc tests were used to compare the data. *, **, and *** denote a significant difference between groups of *P* < 0.05, *P* < 0.01, and *P* < 0.001, respectively. For trends, where 0.05 < *P* < 0.10, p values are reported.

To test the efficacy of shRNA constructs to knock-down *Nfe2l1*, we first tested them in murine embryonic fibroblasts (MEFs). Cells were treated with Bortezomib for 4 hours, which triggers the bounce-back response and causes the cleaved form of Nrf1 (Nrf1 CL) to accumulate in the cytosol (**Figure S5A**) ^27, 32^. The best of the three shRNAs tested efficiently knocked-down cleaved Nrf1 CL and, to a lesser extent, full-length Nrf1 (Nrf1 FL) protein (**Figure S5B**).

Next, we electroporated TA muscle of control and TSCmKO mice with this Nrf1-kd shRNA or scramble shRNA (Scr) as a control. Two weeks post-electroporation, *Nfe2l1* transcripts were more than 60% lower and *Vcp*, along with transcripts coding for all measured 26S proteasome subunit genes, but not the proteasome activator *Psme4*, were significantly reduced by Nrf1-kd shRNA in both control and TSCmKO mice compared to Scr shRNA (**Figure 4B**). Similarly, Nrf1-kd shRNA strongly reduced the protein levels of full-length (FL) Nrf1 in TSCmKO mice and both the cleaved (CL) and alternatively spliced (LCR-F1 or Nrf1β) Nrf1 isoforms along with p97 and OGT in both control and TSCmKO mice compared to a Scr shRNA (**Figure 4C**). On the other hand, Nrf1-kd shRNA significantly increased both the premature and mature forms of SREBF1 in control and TSCmKO mice, suggesting a possible negative feedback loop between Nrf1 and SREBF1. Nrf1-kd shRNA also significantly reduced 26S proteasome protein expression, but unlike at the mRNA level, Nrf1-kd shRNA also moderately reduced PSME4 expression (**Figure 4D**). Furthermore, Nrf1-kd shRNA significantly reduced β1 and β2 and tended to reduce β5 catalytic activity of the 20S proteasome in both control and TSCmKO mice (**Figure 4E**). Despite lower proteasome content and activity, ubiquitinated protein accumulation was lowered by Nrf1-kd shRNA compared to the control Scr shRNA (**Figure 4F)**. However, despite strong reductions in proteasome subunits and activity, knocking down Nrf1 was insufficient to boost skeletal muscle mass in either control or TSCmKO mice (**Figure 4G**), suggesting that the Nrf1-induced upregulation of 26S proteasome content and activity does not contribute directly to mTORC1-induced skeletal muscle atrophy.

Together, these results suggest that Nrf1 coordinately regulates basal proteasome content in healthy skeletal muscle and augments proteasome content in response to sustained mTORC1 activation but is unlikely to be directly responsible for driving atrophy.

### Feedback inhibition of PKB/Akt signaling drives mTORC1-mediated muscle atrophy

As the Nrf1-induced increase in 26S proteasome content and activity does not seem to cause smaller muscles in TSCmKO mice, we next asked whether feedback inhibition of PKB/Akt and subsequent release of FoxO inhibition could be responsible for mTORC1-driven muscle atrophy ^2^. Under atrophic conditions, many atrophy-related genes, including E3 ubiquitin ligases, such as *Fbxo32, Trim63, Fbxo30* and *Fbxo31* as well as the ubiquitin conjugating factor *Ube4b* and the proteasome activator *Psme4* are predominately controlled by FoxO transcription factors ^44, 45^. To test this hypothesis, we crossed TSCmKO mice with AKT-TG mice expressing an active, myristoylated form of PKBα/Akt1, fused to

EGFP and ERT2 in skeletal muscle ^39^. In AKT-TG mice, the PKBα/Akt1-fusion protein is immediately degraded without tamoxifen-induced stabilization via ERT2 binding ^46, 47^.

Control, AKT-TG, TSCmKO and TSCmKO/AKT-TG (TSC-AKT) mice were injected daily with tamoxifen for 5 or 12 consecutive days. In AKT-TG and TSC-AKT mice, tamoxifen induced a rapid increase in body mass after both 5 and 12 days (**Figure 5A**). Pre and post measures of body composition showed that lean mass accretion was primarily responsible for the increase in body mass after 12 days (**Figure 5B**). 5d and 12d AKT-TG experiments were performed separately, normalized to their respective controls and then, based on comparable values and within-group variation, control groups (WT and TSCmKO) from both experiments were pooled for ease of visualization and statistical comparisons. Fast-twitch muscle mass was significantly increased after 5 days and further increased after 12 days of tamoxifen treatment in both AKT-TG and TSC-AKT mice (**Figure 5C**). Endogenous PKB/Akt activation was low in TSCmKO mice and was not changed after 12 days of tamoxifen treatment, but as expected, tamoxifen resulted in the transgenic PKBα/Akt1-GFP fusion protein being highly phosphorylated in AKT-TG and TSC-AKT mice, which also increased phosphorylation of PKB/Akt target PRAS40 **(Figure 5D)**. Tamoxifen also increased phosphorylation of S6 (S240-244 and S235-236) and 4E-BP1 (S65) in AKT-TG mice but did not further increase mTORC1 signaling in TSC-AKT mice compared to TSCmKO mice (**Figure 5D**). On the other hand, significant main effects were observed for both TSC1 KO and PKB/Akt activation in protein synthesis, as determined by the incorporation of puromycin into newly synthesized proteins ^48^ (**Figure 5E**). Consistent with PKB/Akt activation suppressing FoxO, the expression of many ubiquitin-(*Fbxo32, Trim63, Fbxo30, Mdm2, Traf6, Ube4b* and *Ubb*), stress-(*Gadd45a, Gadd34, Nfe2l2, Nqo1, Sesn1*) and autophagy-related (*Sqstm1, Bnip3* and *Ctsl*) genes, the majority of which are known FoxO targets ^45^, were suppressed in TSC-AKT mice (**Figure 5F**). In line with Nrf1 being primarily responsible for mTORC1-mediated proteasome biogenesis, PKB/Akt activation did not alter the majority of 26S proteasome subunits at the transcript (**Figure 5G**) or protein level (**Figure 5H**), with the notable exceptions of *Psmd8* and *Psme4* mRNA. However, despite having a limited impact on proteasome subunit content, prolonged PKB/Akt activation led to an increase in 20S peptidase activity (**Figure 5I**). Together, these data indicate that the FoxO-mediated increase in atrophy-related genes, rather than the Nrf1-induced increase of 26S proteasome content, is responsible for shifting proteostatic balance towards muscle atrophy in TSCmKO mice.

**Figure 5:**
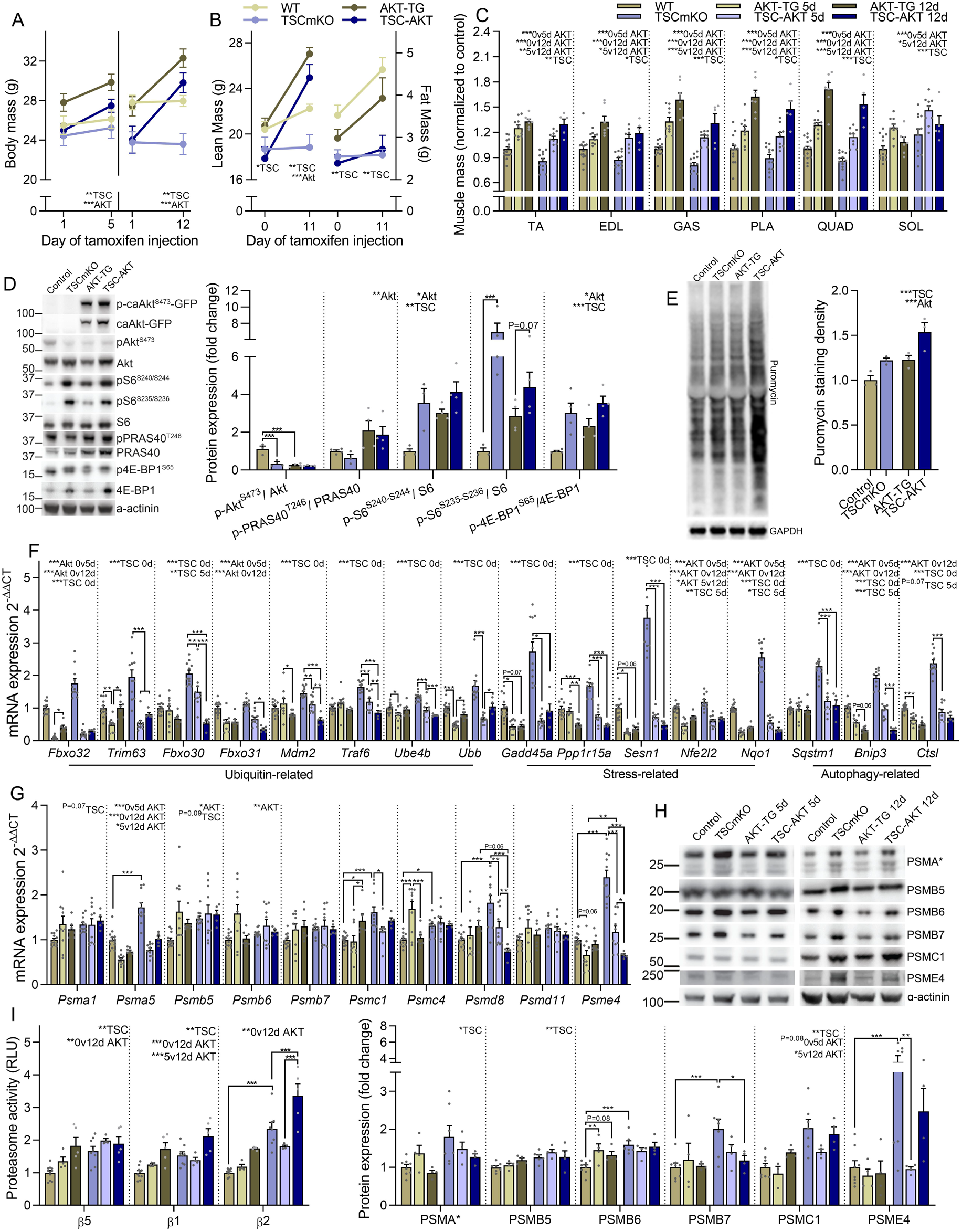
Feedback inhibition of PKB/Akt signaling drives mTORC1-mediated muscle atrophy by upregulating pro-atrophy, FoxO targets. **(A)** body mass before and after 5 or 12 days of tamoxifen treatment (40 mg.kg^-1^.day^-1^) in control and TSCmKO mice and control (AKT-TG) and TSCmKO (TSC-AKT) mice expressing an active, myristoylated form of PKBα/Akt1, fused to EGFP and ERT2 ^39^. Changes in whole-body **(B)** lean (left) and fat (right) mass following 12 days of tamoxifen treatment (n=5-7 per group) and **(C)** muscle mass after 5 and 12 days of tamoxifen treatment in control (n=14), TSCmKO (n=12), AKT-TG (n=10 for 5d and 5 for 12d) and TSC-AKT (n=9 for 5d and 5 for 12d) mice. **(D)** Western blots and quantification of phosphorylated and total proteins involved in the AKT-mTORC1 signaling pathway (n=3-4; *tibialis anterior*) and **(E)** Western blots and quantification of newly synthesized proteins in muscles 30 min after puromycin injection in control, TSCmKO, AKT-TG and TSC-AKT mice treated with tamoxifen for 12 days (n=3; *tibialis anterior*). **(F)** mRNA expression of ubiquitin-stress- and autophagy-related genes induced by sustained mTORC1 activity (*gastrocnemius*), **(G)** mRNA (*gastrocnemius*) and **(H)** protein expression (*tibialis anterior*) of 26S proteasome subunits and the 20S activator PSME4 and **(I)** Luciferase-based peptidase activity (*plantaris*) of 20S proteasome catalytic enzymes in control, TSCmKO, AKT-TG and TSC-AKT mice treated with tamoxifen for 5 or 12 days. For F and G, n=12-14 (Con), 11 (TSCmKO), 10 (5d AKT-TG and TSC-AKT), 6-7 (12dATK-TG) and 5 (12dTSC-AKT). For H, n=8 (Con), 7 (TSCmKO) and 4 (5d and 12d AKT-TG and TSC-AKT). For I, n=10 (Con), 8 (TSCmKO) and 4 (5d and 12d AKT-TG and TSC-AKT). *Tubb* was used as the reference gene for F and G, while α-actinin (D and H) or GAPDH (E) were used as the protein loading control. Data are presented as mean ± SEM. Two-way ANOVAs with Sidak post hoc tests were used to compare the data. *, **, and *** denote a significant difference between groups of *P* < 0.05, *P* < 0.01, and *P* < 0.001, respectively. For trends, where 0.05 < *P* < 0.10, p values are reported.

### PKB/Akt reactivation compromises muscle integrity in TSCmKO mice

Although restoration of PKB/Akt activity in TSCmKO mice blocks atrophy-related gene expression and potently induces muscle growth, while examining the muscle structure of TSC-AKT mice we observed a striking accumulation of aberrant muscle fibers containing multiple vacuole-like structures **(Figure 6A)**. Since PKB/Akt blocks UPS induction, we hypothesized that damaged proteins that would normally be degraded by the UPS may be directed towards the autophagy/lysosomal system. However, since autophagy is blocked by sustained mTORC1 activity, protein aggregates marked for breakdown by the autophagy receptor p62 would accumulate. Indeed, strong p62 staining was observed in TA muscle fibers from TSC-AKT, but not control, TSCmKO or AKT-TG mice **(Figure 6B)**. Many fibers strongly positive for p62 staining also contained unstained regions indicative of the vacuoles observed in H&E stains. Importantly, p62 accumulation and vacuolated fibers are not normally seen in young TSCmKO mice and are rather characteristic of the late-onset myopathy typically observed at an age of 9 to 12 months ^4^. This indicates that the mTORC1-driven increase in the expression of atrogenes regulated by FoxO transcription factors is a protective response that can compensate, at least initially, for sustained autophagy inhibition. We next wondered whether the Nrf1-mediated increase in proteasome content driven by mTORC1 also plays a similar role. To this end, we looked for signs of disturbed proteostasis in muscle from control-Scr, Con-Nrf1-kd, TSCmKO-Scr and TSC-Nrf1-kd mice. While Nrf1 depletion significantly suppressed some ubiquitin-stress-and autophagy-related genes, in both control and TSCmKO mice, it also increased the expression of key ubiquitin-related (*Fbxo32, Trim63, Eif4ebp1, Gadd45a, Ppp1r15a*), stress-related (*Nqo1*) and autophagy-related (*Sqstm1* and *Bnip3*) genes specifically in TSCmKO muscle **(Figure 6C and D)**. Furthermore, a significant accumulation of BNIP3 and p62 protein was observed in TSCmKO-Nrfkd1 compared to TSCmKO muscle **(Figure 6E)**, indicating an impaired capacity to degrade damaged proteins. Together, these data point to seemingly conflicting roles of UPS induction in the response to sustained mTORC1 activity, on the one hand promoting muscle atrophy, while on the other hand compensating for autophagy blockade and preserving muscle integrity.

**Figure 6:**
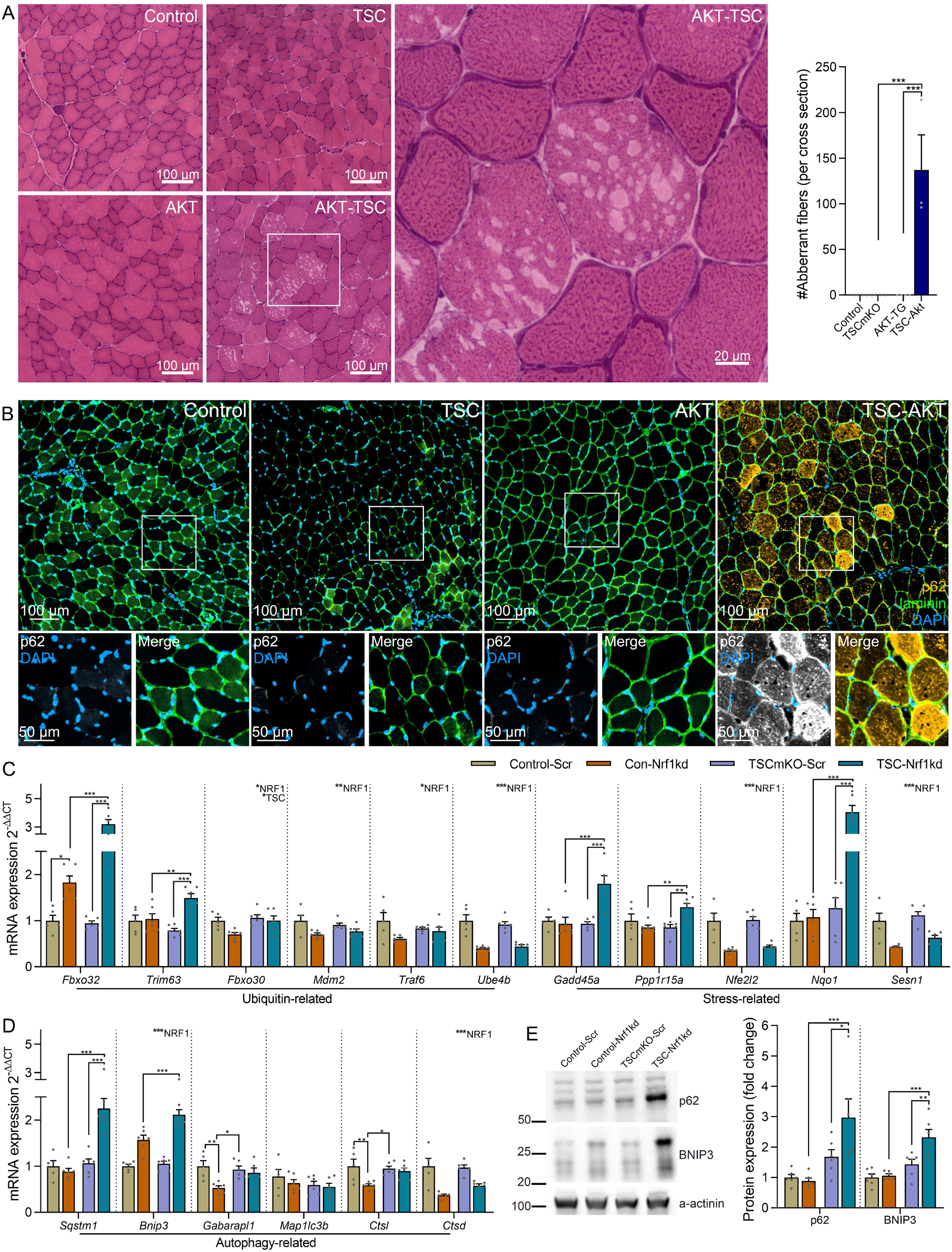
mTORC1-induced activation of the ubiquitin proteasome system preserves muscle integrity. **(A)** Representative images of Hematoxylin and eosin-stained cross sections and quantification of aberrant fibers and **(B)** representative images of *tibialis anterior* cross sections stained for p62 (yellow), laminin (green) and DAPI (blue) in control, TSCmKO (TSC), AKT-TG (AKT) and TSC-AKT mice after 12 days of tamoxifen treatment. **(C)** mRNA expression of ubiquitin-stress- and **(D)** autophagy-related genes in *gastrocnemius* muscle and **(E)** Western blots and quantification of autophagy-related protein expression in *tibialis anterior* muscle of control and TSCmKO mice 2 weeks after transfection of a plasmid expressing either a scrambled (Scr) or *Nfe2l1*-targeting (Nrf1kd) shRNA. *Des* was used as the reference gene and α-actinin as the protein loading control. For all experiment n=6 per group, except for control in C and D where n=5. Data are presented as mean ± SEM. Two-way ANOVAs with Sidak post hoc tests were used to compare the data. *, **, and *** denote a significant difference between groups of *P* < 0.05, *P* < 0.01, and *P* < 0.001, respectively. For trends, where 0.05 < *P* < 0.10, p values are reported.

## Discussion

Skeletal muscle is highly dynamic, adapting its size to meet the specific needs of the body, with increased loading driving hypertrophy and disuse causing atrophy. An interconnected network of processes and signaling pathways act in unison to regulate changes in muscle size while maintaining the health of the proteome. Seminal papers place the PKB/Akt-mTORC1-FoxO pathway at the heart of this network ^49, 50, 51, 52^ and led to the model that muscle size is determined by the balance between protein synthesis and degradation. According to this model, muscle mTORC1 activation, which increases protein synthesis and inhibits autophagy, should drive hypertrophy and block atrophy. However, this assumption does not hold true, with neither short-(**Figure 2**) or long-term ^53^ mTORC1 activation through muscle fiber-specific *Tsc1* deletion resulting in hypertrophy. On the contrary, chronic mTORC1 activation reduces fast-twitch fiber size and the mass of all fast-type muscles. Moreover, TSCmKO mice show many features consistent with accelerated muscle aging ^6, 54^. While the sarcopenia-like myopathy takes ∼9 months to develop in TSCmKO mice ^4, 6^, with multiple lines of evidence implicating impaired autophagy as the mediating factor, mTORC1-driven muscle atrophy is already seen in 3-month-old

TSCmKO mice ^53^ and after just 21d of tamoxifen-induced *Tsc1* KO (**Figure 2**). Here, we investigated the mechanisms underlying mTORC1-driven muscle atrophy in young, pre-myopathic TSCmKO mice and following acute TSC1 depletion in young mice, thereby avoiding influence from secondary consequences of the late-onset myopathy. Using genetic tools to activate and uncouple mTORC1 and PKB/Akt from potential upstream influences, chemical inhibition of proteasome activity and shRNA-mediated proteasome depletion, as well as transcriptomic and proteomic profiling, we discover that an mTORC1-driven upregulation of the ubiquitin proteasome system, including Nrf1-mediated proteasome biogenesis, outstrips increased translation and explains the counterintuitive atrophy in TSCmKO mice. Furthermore, we uncover seemingly antagonistic roles of mTORC1-driven UPS activation in regulating muscle proteostasis, promoting atrophy while simultaneously maintaining muscle integrity.

UPS activation is a common feature of many muscle wasting conditions and loss and gain of specific E3 ligase function, such as atrogin1 ^55^ blunts and promotes muscle atrophy, respectively. As such, perturbing UPS activity has received considerable attention as a potential means of tackling various muscle wasting conditions. Here we provide substantial evidence that hyperactive mTORC1 activates the UPS, including the induction of many FoxO-regulated pro-atrophy E3 ligases through PKB/Akt suppression and ubiquitous upregulation of proteasome subunits via Nrf1, in fast but not slow twitch muscles accounting for their atrophic and hypertrophic responses to mTORC1 activation, respectively. Interestingly, a similar increase in expression of atrogenes and proteasome subunit genes was also observed in muscle from naturally aged mice, in which mTORC1 activity is high, and whose function and size is improved by prolonged rapamycin treatment ^6^.

Nrf1 transcriptionally promotes proteasome biogenesis by binding ARE sequences within the promoter region of all proteasome subunit genes, and its induction has been described, predominately in cultured cells, in response to proteasome inhibition ^43^ and mTORC1 activity through SREBF1 ^35^. Here, we describe for the first time a role for Nrf1-mediated proteasome biogenesis in response to sustained mTORC1 activity in skeletal muscle. While co-activation of UPS-mediated breakdown alongside protein synthesis by mTORC1 may seem like an exercise in futility, our data suggests this may in fact represent a prudent strategy to cope with the inherent errors associated with protein translation, processing and folding ^56^. That is, mTORC1 may stimulate proteasome biogenesis to reduce the risk of proteotoxicity resulting from a build-up of damaged and misfolded proteins created by higher rates of protein synthesis. In support of this notion, UPS impairment either through PKB/Akt activation or proteasome depletion resulted in signs of proteotoxicity in TSCmKO mice with sustained mTORC1 activity. Misfolded proteins are predominately tagged with K48-linked ubiquitin chains and therefore preferentially degraded by the UPS. However, if the UPS is overwhelmed, ubiquitinated misfolded proteins can aggregate and form condensates through p62-mediated phase separation ^57^. Since free mono-ubiquitin released by proteasomes during the de-ubiquitination step impairs p62-mediated phase separation, condensate formation is blocked when proteasome activity is appropriate ^57^. If UPS activity is insufficient, p62-positive condensates would normally form and then be degraded through selective autophagy. We have previously observed widespread p62-positive condensate staining and vacuoles in muscle from TSCmKO mice ^4^. However, these features do not become prevalent until 6-9 months of age, suggesting that Nrf1-mediated upregulation of the UPS by mTORC1 is initially sufficient to prevent a buildup of damaged/misfolded proteins and subsequent p62-positive condensate formation. In line with this, rapid p62 accumulation was observed within two weeks of UPS abrogation by either Nrf1-kd-mediated proteasome depletion or PKB/Akt-FoxO-dependent suppression of UPS induction **(Figure 6)**. However, the UPS cannot completely replace autophagic breakdown, since it is structurally limited to the breakdown of unfolded proteins that fit within the core particle ^9^. The late-onset accumulation of p62-positive condensates and vacuoles in TSCmKO mice could thus result from the accumulation of damaged organelles and other substrates that cannot physically be degraded by the UPS. Similarly, our data provides physiological context for the feedback inhibition from S6K1 to IRS1 on PKB/Akt that is activated by sustained mTORC1 activity. While this negative feedback loop blunts muscle growth we show that it also serves to relieve the proteostatic burden of PKB/Akt activity and maintain muscle integrity. Indeed, chronic PKB/Akt activation has also been shown to disturb muscle proteostasis, with six months of constitutively active PKB/Akt expression also promoting p62-positive aggregates and vacuolated fibers along with other myopathic features ^58^.

While knockdown of Nrf1 and depletion of proteasomes in skeletal muscle led to increased signs of damaged protein accumulation, including increases in p62 and BNIP3, as well as induction of ubiquitin-(*Fbxo32* and *Trim63*) and stress-(*Gadd45a, Ppp1r15a* and *Nqo1*) related atrogenes, the majority of which are FoxO targets, in TSCmKO muscle, Nrf1-kd was insufficient to rescue mTORC1-induced muscle atrophy. Therefore, it would seem that increased proteasome content alone is not sufficient to initiate proteasome degradation and therefore muscle atrophy; rather, the presence of FoxO-regulated enzymes involved in ubiquitination appears to be the limiting factor controlling muscle atrophy. Consistently, inhibition of one ubiquitin system gene, such as *Fbxo32, Trim63, Fbxo30* or *Traf6* is sufficient to limit protein degradation and attenuate muscle loss in various atrophic conditions ^13, 55, 59^. In line with these observations, PKB/Akt activation strongly suppressed ubiquitin-related FoxO target genes and potently stimulated muscle hypertrophy in both control and TSCmKO mice (**Figure 5**).

In a perfect world, only misfolded, damaged and obsolete signaling proteins would be targeted for destruction by the proteasome, however, the fact that overexpression of specific E3 ligases such as atrogin1 ^55^ is sufficient to drive muscle wasting in otherwise healthy muscle tissue means that functional proteins are also caught in the cross hairs of FoxO-mediated UPS activation. A similar phenomenon appears to occur as a result of mTORC1-driven UPS induction, since 1) PKB/Akt-activation drives rapid muscle hypertrophy in TSCmKO mice and 2) the slow-twitch soleus muscle, which fails to induce E3 ligase gene expression (e.g. *Fbxo32* and *Trim63*) despite PKB/Akt suppression, also displays strong hypertrophy in TSCmKO mice. However, it is important to note that the mass gained by the SOL muscle in TSCmKO mice is largely non-functional, since muscle force is severely perturbed, as in fast-twitch muscle fibers ^4^.

Together, these data show that mTORC1-induced activation of the UPS has dual roles in muscle proteostasis, contributing to muscle atrophy through PKB/Akt suppression and FoxO-mediated induction of atrogenes, but simultaneously preserving muscle integrity by degrading damaged and misfolded proteins (**Figure 7**). Since mTORC1 activation and autophagy impairment are frequently observed alongside UPS activation in atrophic conditions such as sarcopenia and denervation ^4, 6^, our data indicate that great caution should be exercised with intervention strategies designed to blunt the UPS as a means of reducing muscle atrophy. Furthermore, the evaluation of such intervention strategies should include measures of both muscle size and proteome health.

**Figure 7:**
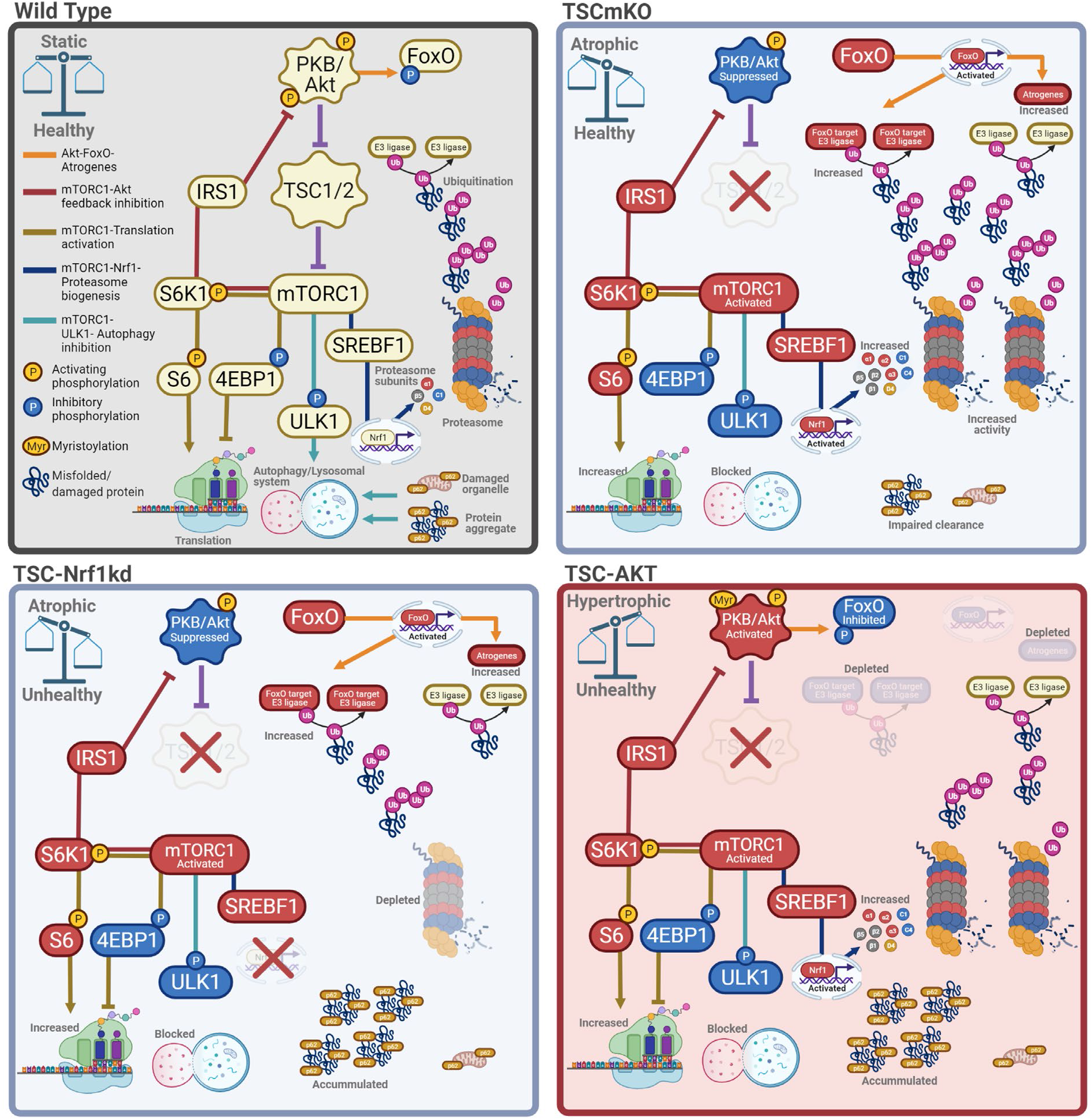
Proteostatic consequences of PKB/Akt-mTORC1 pathway manipulation in muscle. Major signaling networks involved in PKB/Akt-mTORC1-mediated muscle proteostasis in control mice (wild type, upper left), with sustained mTORC1 activity (TSCmKO, upper right), sustained mTORC1 activity and Nrf1-kd-mediated proteasome depletion (lower left) and sustained mTORC1 activity with constitutive PKB/Akt activation (lower right). Impairing ubiquitin-proteasome-mediated protein breakdown under conditions with high mTORC1 activity disrupts proteostasis and leads to unhealthy muscles, independent of muscle growth status. Figure created with BioRender.com.

## Materials and methods

### Animal care

All procedures were performed in accordance with Swiss regulations for animal experimentation, and approved by the veterinary commission of the Canton Basel-Stadt. TSCmKO mice and their genotyping were previously described ^3^. Controls were littermates floxed for *Tsc1* but not expressing HSA-Cre recombinase. Rapamycin (8 mg/kg; LC Laboratories) was administered I.P. as previously described ^3^ for three consecutive days. Bortezomib (1mg/kg; Selleck Chemicals) or saline was administered I.P. every 72 hours for four weeks and muscles were collected 6 h following the final injection. *Tsc1*-floxed mice expressing a tamoxifen inducible HSA-Cre (iTSCmKO) and their genotyping were previously described ^39^. Littermates floxed for *Tsc1* but not expressing Cre recombinase were used as controls. To induce ablation of *Tsc1* in Cre recombinase expressing muscle fibers, tamoxifen (Sigma-Aldrich) diluted in corn oil (Sigma-Aldrich) was administered I.P. at a dose of 75 mg·kg·day^-1^ for five consecutive days. The day after the fifth injection was defined as day 1 and Tamoxifen was administered again on days 3 and 4 (**Figure S3A**). Both iTSCmKO mice and their controls received tamoxifen. AKT-TG mice expressing a tamoxifen-inducible constitutively active form of PKB/Akt were obtained from Dr. David Glass at Novartis Institutes for BioMedical Research (NIBR, Cambridge, MA, USA). Generation and genotyping of AKT-TG mice were previously described ^39^. TSC-AKT mice were obtained by crossing *Tsc1*-floxed mice expressing HSA-Cre with AKT-TG mice. Control mice for TSC-AKT and AKT-TG mice were littermates floxed for *Tsc1* but not expressing Cre recombinase or the Akt transgene. AKT-TG, TSC-AKT mice and their respective controls were administered tamoxifen (40 mg·kg·day^-1^) I.P. for 5 or 12 days. Body composition measurements were performed using an EchoMRI-100 (EchoMRI Medical Systems). Tissue was collected within 6 h after final tamoxifen injection on the 5^th^ or 12^th^ treatment day. For protein synthesis experiments, puromycin (0.04 μmol·g^-1^; Sigma P8833) was administered I.P. exactly 30 min before dissection. Experiments were performed on 3-4 month old female and male mice, except for those using tamoxifen where only male mice were used. Age-matched mice of the same sex were used for each individual experiment. Mice were maintained in a licensed animal facility with a fixed 12 h light-dark cycle (23°C) and were allowed free access to food and water.

### Cell culture

SV40 immortalized mouse embryonic fibroblasts (MEFs) were maintained in Glutamax Dulbecco’s modified Eagle’s medium (DMEM Glutamax and pyruvate, Gibco) supplemented with 10% fetal bovine serum (FBS, Gibco) and 1% penicillin-streptomycin (pen/strep, Gibco) on cell culture dishes in 37°C incubator with 5% CO2. For Nrf1-kd experiment, MEFs were transfected with plasmids encoding for Nrf1-kd shRNA (see below) using Lipofectamine 2000 (Invitrogen). 48 hours after transfection cells were scraped and lysed in ice cold RIPA buffer (50 mM TrisHCl pH 8.0, 150 mM NaCl, 1% NP-40, 0.5% sodium deoxycholate, 0.1% SDS, ddH2O) supplemented with phosphatase and protease inhibitors (Roche). DNA was sheared with a syringe (26G needle). Afterwards, the lysate was centrifuged at 16,000 g for 20 min at 4°C. Supernatant (cleared lysates) were used to determine total protein amount using the Pierce BCA Protein Assay Kit (Thermo Fisher Scientific) according to manufacturer’s protocol. For Nrf1 accumulation experiment, MEFs were treated with bortezomib (BTZ; 10 nM, Selleck Chemicals) or vehicle (DMSO; Sigma-Aldrich) for four hours prior to lysis.

### RNA extraction for RNA-seq

EDL muscles isolated from control and TSCmKO mice, were used to generate the corresponding cDNA-libraries. Poly-A mRNA was directly isolated from frozen EDL samples by using the Dynabeads™ mRNA DIRECT™ Purification Kit (61011, Invitrogen) with the Mini configuration followed by alkaline hydrolysis. To provide 5’ to 3’ directionality to fragmented samples, mRNA was treated with phosphatase and then with polynucleotide kinase (PNK). Further, 3’ and 5’ adaptors (Illumina) were ligated and the resulting product was reverse transcribed to generate cDNA by PCR. During PCR amplification, each prepared sample was uniquely indexed (barcoding) using index primer (Illumina) for multiplexing (differently indexed samples in one lane). PCR products have been purified twice with AMPure beads (Agencourt). The quality of the cDNA library was verified and quantified with a Bioanalyzer (Agilent Technologies). Illumina “HiSeq 2000” was used for sequencing at the Quantitative Genomics Facility (QGF) at the Department of Biosystems Science and Engineering (D-BSSE) of the ETH Zürich in Basel.

### Protein isolation for Tandem Mass Tag (TMT)-LC-MS/MS

Dissected muscle was rapidly frozen in liquid nitrogen. Frozen muscles were pulverized on a metal plate chilled in liquid nitrogen and directly snap-frozen as a pellet in liquid nitrogen. Samples were lysed in ice cold lysis buffer (1% sodium deoxycholate, 100 mM ammoniumbicarbonate) and ultra-sonicated twice for 10 s with a VialTweeter (Hielscher). Afterwards, the lysate was centrifuged at 16,000 g for 10 min at 4°C. Supernatant (cleared lysates) were used to determine total protein amount using the Pierce BCA Protein Assay Kit (Thermo Fisher Scientific) according to manufacturer’s protocol. Next, protein was reduced and alkylated. Afterwards, proteins were digested using Trypsin digestion at 37°C overnight. Samples were purified (solid phase extraction), labeled with TMT10plex Label Reagents (Thermo scientific) and then mixed before sample fractionation and clean-up. Labeled samples are analyzed by high resolution Orbitrap LC-MS/MS.

### Data Processing

RNA-seq data were processed as previously described and have been previously reported ^6^ and deposited at the Gene Expression Omnibus GEO) ^60^ under accession number GSE139214. Additionally, the data are freely available using the web-based application, SarcoAtlas (https://sarcoatlas.scicore.unibas.ch/). Mass-spectrometric data was statistically validated by the SafeQuant software tool ^61^ developed in house. Proteins were considered as significantly differentially expressed with an adjusted p value of 0.05. Characterization and enrichment analysis of the differentially expressed proteins was done using DAVID analysis and functional annotation clustering ^12, 62^.

### RT-qPCR

Dissected muscle was rapidly frozen in liquid nitrogen. Total RNA was extracted using the RNeasy Mini Kit (Qiagen) according to manufacturer’s protocol. Equal amounts of RNA were transcribed into cDNA using the iScript cDNA Synthesis Kit (BioRad). Selected genes were amplified and detected using the Power SYBR Green PCR Master Mix (Applied Biosystems) or FastStart Essential DNA Green Master (Roche). Quantitative expression was determined by StepOnePlus Real-Time PCR System (Applied Biosystems) or LightCycler 480 (Roche). Data were analyzed using the comparative Cq method (2−ΔΔCq). Raw Cq values of target genes were normalized to Cq values of a housekeeping gene (*Tubb, Actb* or *Des*), which was stable between conditions, and then further normalized to the control group for ease of visualization. Primers used are outlined in **Table S2**.

### In vitro muscle force

*In vitro* muscle force was measured in EDL muscles carefully excised and mounted on the 1200 A Isolated Muscle System (Aurora Scientific, Aurora, ON, Canada) in an organ bath containing 60 mL of Ringer solution (137mM NaCl, 24mM NaHCO3, 11mM glucose, 5mM KCl, 2mM CaCl2, 1mM MgSO4, and 1mM NaH2PO4) gassed with 95% O2, 5% CO2 at 30 °C. After defining the optimal length, muscles were stimulated with 15-V pulses. Muscle force was recorded in response to 500-ms pulses at 10–250 Hz.

### Western blot analysis

Dissected muscle was rapidly frozen in liquid nitrogen. Frozen muscles were pulverized on a metal plate chilled in liquid nitrogen and directly snap-frozen as a pellet in liquid nitrogen. Samples were lysed in ice cold RIPA buffer (50 mM TrisHCl pH 8.0, 150 mM NaCl, 1% NP-40, 0.5% sodium deoxycholate, 0.1% SDS, ddH2O) supplemented with phosphatase and protease inhibitors (Roche), incubated on a rotating wheel for 2 h at 4°C and sonicated twice for 10 s. Afterwards, the lysate was centrifuged at 16,000 g for 20 min at 4°C. Supernatant (cleared lysates) were used to determine total protein amount using the Pierce BCA Protein Assay Kit (Thermo Fisher Scientific) according to manufacturer’s protocol. Proteins were separated on 4-12% Bis-Tris Protein Gels (NuPage Novex, Thermo Fisher Scientific) and transferred to nitrocellulose membrane (GE Healthcare Life Sciences, Amersham). The membrane was blocked with 5% BSA, 0.1% Tween-20, PBS for 1 h at room temperature. The primary antibody diluted in the blocking solution was incubated overnight at 4°C with continuous shaking. The membranes were washed with PBS-T (0.1% Tween-20, PBS) for 7 minutes three times and incubated with secondary horseradish peroxidase-conjugated (HRP) antibody for 1 h at room temperature. After washing with PBS-T, proteins were visualized by chemiluminescence (KPL LumiGLO®, Seracare). Signal was captured on a Fusion Fx machine (VilberLourmat) and analyzed with FUSION Capt FX software. All antibodies used for immunoblotting are listed in **Table S3**.

### Proteasome activity assay

Proteasome activity measurements were adapted from a protocol described previously (Strucksberg et al., 2010). Briefly, dissected *extensor digitorum longus* (EDL) or *plantaris* (PLA) muscles were rinsed in ice-cold PBS, immediately cut into 5-6 pieces and directly lysed in ice-cold PBS-E (5 mM EDTA pH 8.0, PBS pH 7.2) and sonicated two times for 10 s. Afterwards, the lysate was centrifuged at 13,000 g for 5 min at 4°C. Supernatant (cleared lysates) were used to determine total protein amount using the Pierce BCA Protein Assay Kit (Thermo Fisher Scientific) according to manufacturer’s protocol. Three individual luciferase-based Proteasome-Glo™ Assay Systems (Promega) were used to measure the activity of each peptidase of the proteasome. Assay was performed on white, 96-well microplates (greiner BIO-ONE) and luminescence was measured with an Infinite M1000 (Tecan). Human, purified 20S proteasome (Enzo Life Sciences, BML-PW8720) was used as a positive control. Proteasome inhibitor MG-132 (50 µM, TOCRIS Biotechne) was used to subtract non-proteasomal background activity.

### Histology analysis

Cryostat sections (10 μm) were cut from TA as previously described ^6^. For histological analysis TA sections were stained with hematoxylin and eosin (H&E; Merck, Zug, Switzerland). To determinate fiber type and fiber size sections were blocked and permeabilized in PBS containing 10% goat serum and 0.4% triton X-100 for 30 min before being incubated for 2 h at RT in a primary antibody solution containing BA-D5, SC-71, BF-F3 and laminin (#L9393, Sigma), and 10% goat serum. BF-F3, BA-D5, and SC-71 antibodies were developed by Prof.Stefano Schiaffino and obtained from the Developmental Studies Hybridoma Bank developed under the auspices of the National Institute of Child Health and Human Development, and maintained by the University of Iowa Department of Biology. Sections were washed four times for 10 min in PBS and then incubated in a secondary antibody solution containing DyLight 405 (#115-475-207, Jackson), Alexa568 (#A-21124, Invitrogen), Alexa488 (#A-21042, Invitrogen), Alexa647(#711-605-152, Jackson), and 10% goat serum. Sections were then washed four times for 10 min in PBS and mounted with Vectashield Antifade Mounting Medium (Vectorlabs). For p62 staining, sections were fixed in 4% PFA for 10 min prior to immunostaining. Primary antibodies used were p62/SQSTM1 (GP62-C, Progene) and Laminin-α2 (#ab11576, Abcam). Secondary antibodies were Alexa488 (#112-545-003, Jackson) and Cy3 (#706-165-148, Jackson). Muscle sections were imaged at the Biozentrum Imaging Core Facility with an Axio Scan.Z1 Slide Scanner (Zeiss-Oberkochen, Germany) equipped with appropriate band-pass filters. Fiji macros were developed in-house to allow an automated analysis of muscle fiber types (based on intensity thresholds) and muscle cross-sectional area (i.e., minimal Feret’s diameter, based on cell segmentation). All macros and scripts used in this study are available upon request.

### RNAscope

Slides were fixed in cold 4% PFA for 15 min at 4 °C before serial dehydration for 5 min in each of 50%, 70% and 2 × 100% ethanol. Slides were then dried for five min at RT and circled with a Hydrophobic Barrier Pen (Vector Laboratories, GZ-93951-68). Sections underwent protein digestion (protease IV) for 30min and then 15min at RT before being washed twice with PBS. RNA hybridization with probes against *Nfe2l1* (580611, Advanced Cell Diagnostics) and subsequent amplification steps were performed according to the manufacturers instructions at 40°C in a HybEZ™ oven (Advanced Cell Diagnostics). After hybridization, Slides were blocked 60 min at RT in PBS containing 0.4% Trition X-100 and 10% goat serum, washed 2 × 5 min in PBS and then incubated with primary antibodies against Laminin-α2 (#11576, Abcam) in antibody solution containing 10% goat serum in PBS. Slides were then washed 4 × 10 min in PBS and incubated with GARt-488 (112-545-008, Jackson) secondary antibody and DAPI in antibody solution. Slides were then washed 4 × 10 min in PBS and mounted with ProLong™ Gold antifade (Invitrogen).

### In vivo muscle transfection and electroporation

The methods to construct plasmid vectors encoding shRNA have been described elsewhere ^63^. The murine 19 nucleotide target sequences corresponding to: GGC CCG ATT GCT TCG AGA A (Nrf1) and pRFP-C-RS scrambled shRNA plasmid vectors were obtained from OriGene (TR30015). For analgesia, mice were administered Buprenorphine (0.1 mg/kg) subcutaneously before and every 4 to 6 hours during the day and in drinking water overnight for 2 days and nights after electroporation. Mice were anesthetized with isoflurane. The *tibialis anterior* (TA) muscle was exposed and injected with 50 μl (8 IU) of hyaluronidase (Sigma) two hours before being injected with 50 μl of 2 μg/μl shRNA plasmid DNA. Electroporation was then performed by applying three 30 ms pulses of 150 V/cm with a 50 ms pulse interval using a NEPA21 electroporation system (NepaGene). Mice were analyzed 2 weeks after electroporation.

### Statistical analysis

All values are expressed as mean ± SEM, unless otherwise stated. Data were tested for normality and homogeneity of variance using a Shapiro–Wilk and Levene’s test, respectively. Data were analyzed in GraphPad Prism 8. Student’s t tests were used for pairwise comparisons, while one-way ANOVAs with Fisher’s LSD post hoc tests were used to compare between three groups, so long as the ANOVA reached statistical significance. Two-way ANOVAs with Sidak post hoc tests, or two-way repeated-measure ANOVAs for multiple recordings over time, were used to compare between groups with two independent variables. Both significant differences (P < 0.05) and trends (P < 0.1) are reported where appropriate.

## Acknowledgments

We thank the Biozentrum In-house Imaging Core Facility, Proteomics Core Facility and the Quantitative Genomics Facility (QGF) at the Department of Biosystems Science and Engineering (D-BSSE, ETH Zürich, Basel) for their technical support.

## Author contributions

MK and GM designed and performed experiments, analyzed data and wrote the manuscript with input from all authors. SL performed muscle force measurements and electroporations. FO performed intraperitoneal injections, behavioral experiments and amplification of plasmid DNA. KC performed behavioral experiments with iTSCmKO mice. LT and NM created cDNA libraries for RNA-sequencing. CZ created proteomics data. DJG provided the AKT-TG mice. MZ supported and supervised the generation of sequencing data and the analysis of RNA-seq data. DJH performed experiments, analyzed data, prepared figures and wrote the paper. MAR conceived the project, secured funding and wrote the manuscript.

