## Supplementary material for "Dual roles of mTORC1-dependent activation of the ubiquitin-proteasome system in muscle proteostasis"

### SUPPLEMENTARY FIGURES

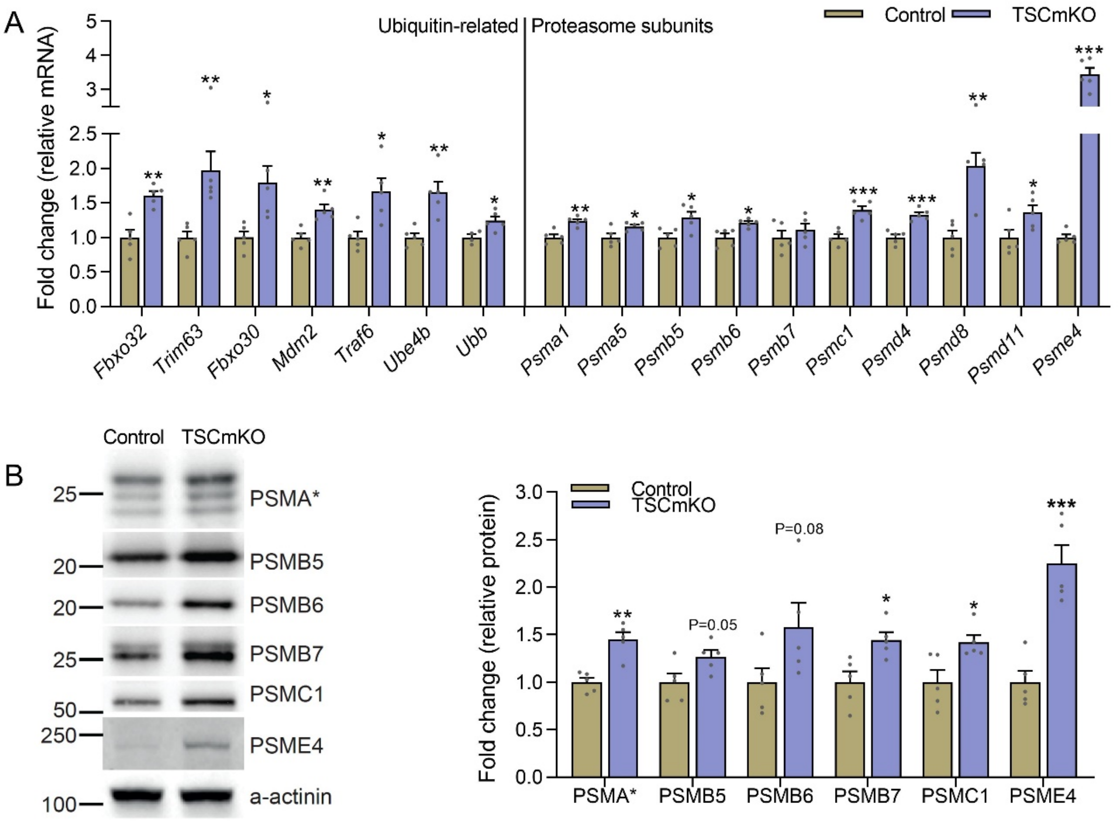


**Figure S1: (A)** mRNA expression of ubiquitin-related (left) and proteasome subunit (right) genes measured by RT-qPCR and **(B)** western blots and quantification of protein expression of 26S proteasome subunits and the 20S activator PSME4 in *gastrocnemius* (mRNA) and *tibialis anterior* (protein) muscle from control and TSCmKO mice. Data are presented as mean ± SEM. Two-tailed Student's t-tests were used to compare the data. *, **, and *** denote a significant difference between groups of $P < 0.05$, $P < 0.01$, and $P < 0.001$, respectively. For trends, where $0.05 < P < 0.10$, $P$ values are reported.

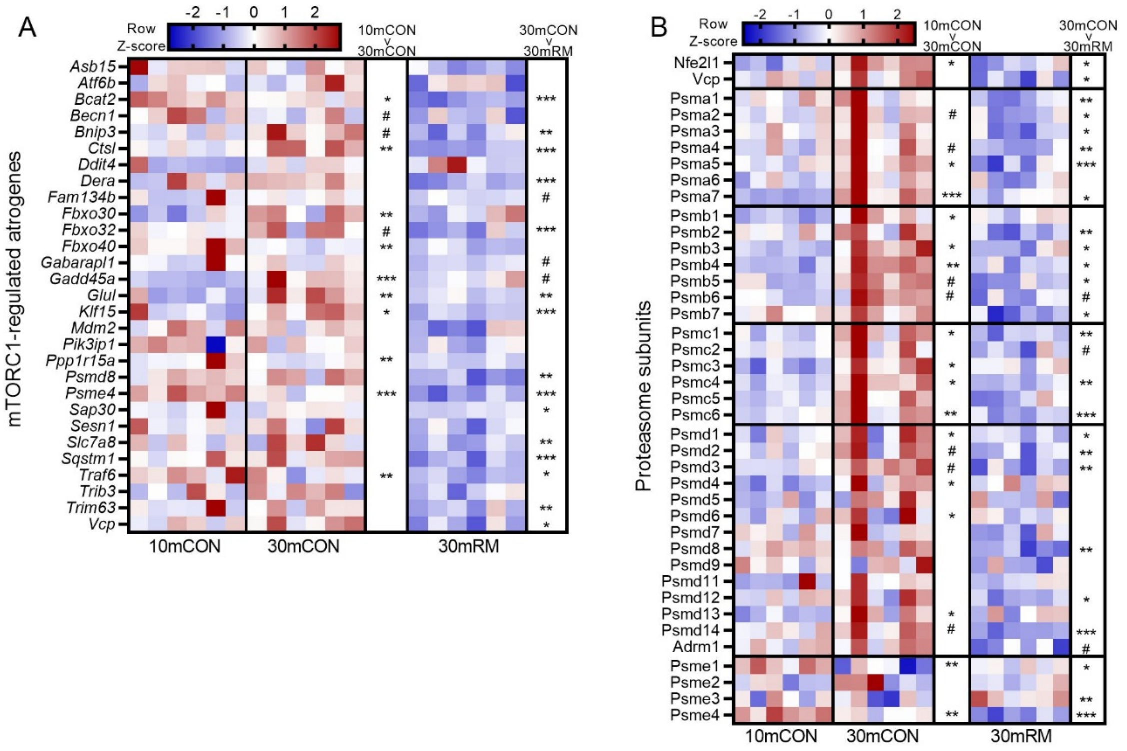


**Figure S2:** Heatmap of changes in mRNA expression of **(A)** mTORC1-regulated atrogenes and **(B)** 26S proteasome subunits and members of the PA28 and PA200 proteasome activators in wild-type adult mice (10mCON), 30-month-old, sarcopenic mice (30mCON) and 30-month-old mice treated from 15 months of age with rapamycin (30mRM) [6]. *, **, and *** denote a significant difference between groups of $P < 0.05$, $P < 0.01$, and $P < 0.001$, respectively. # denotes a trend, where $0.05 < P < 0.10$.

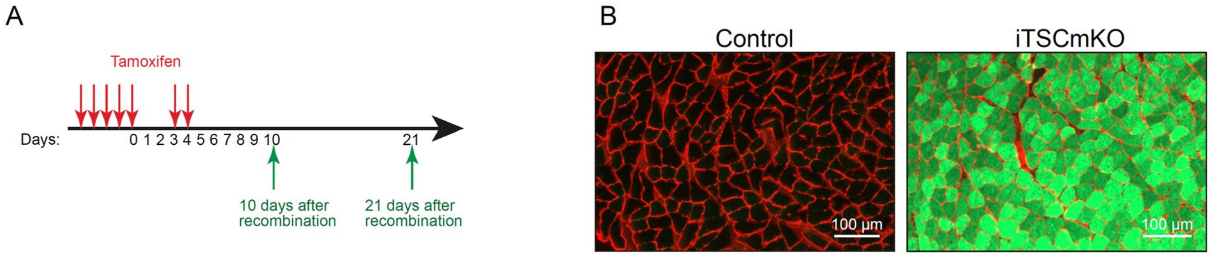


**Figure S3. (A)** tamoxifen treatment schedule for 10 day and 21 day induced TSC1 knockout mice (iTSCmKO). **(B)** Cross sections of *tibialis* anterior muscle from control and 10d iTSCmKO mice that also contain a *Rosa26* knock-in construct expressing enhanced green fluorescent protein (EGFP) as a Cre reporter. While control mice without HSA-MerCreMer do not express EGFP, all fibers of mice with Cre are EGFP positive. Muscle cross sections were also stained with antibodies against laminin (red).

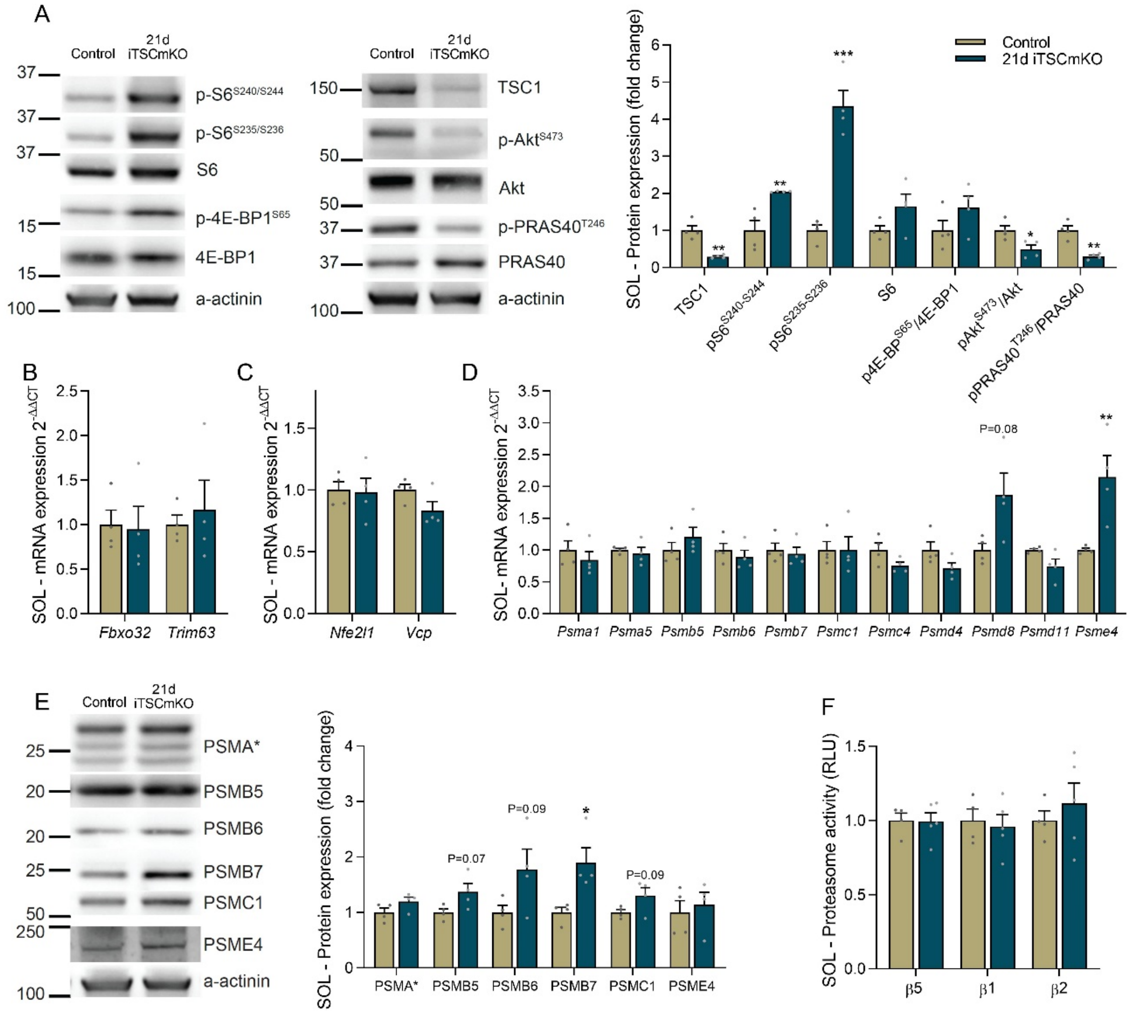


**Figure S4. (A)** Western blots and quantification of phosphorylated and total proteins involved in the PKB/Akt-mTORC1 signaling pathway in *soleus* muscle 21 days after *Tsc1* deletion. mRNA expression of **(B)** atrogenes, **(C)** *Nfe2l1* and related Nrf1 regulators and **(D)** 26S proteasome subunits and the 20S activator *Psme4* in *soleus* muscle 21 days after tamoxifen treatment. **(E)** Western blots and quantification of 26S proteasome subunits and PSME4 and **(F)** luciferase-based peptidase activity of 20S proteasome catalytic enzymes in *soleus* muscle 21 days post-tamoxifen. Data are presented as mean ± SEM. Two-tailed Student's t-tests were used to compare the data. *, **, and *** denote a significant difference between groups of $P < 0.05$, $P < 0.01$, and $P < 0.001$, respectively. For trends, where $0.05 < P < 0.10$, $P$ values are reported.

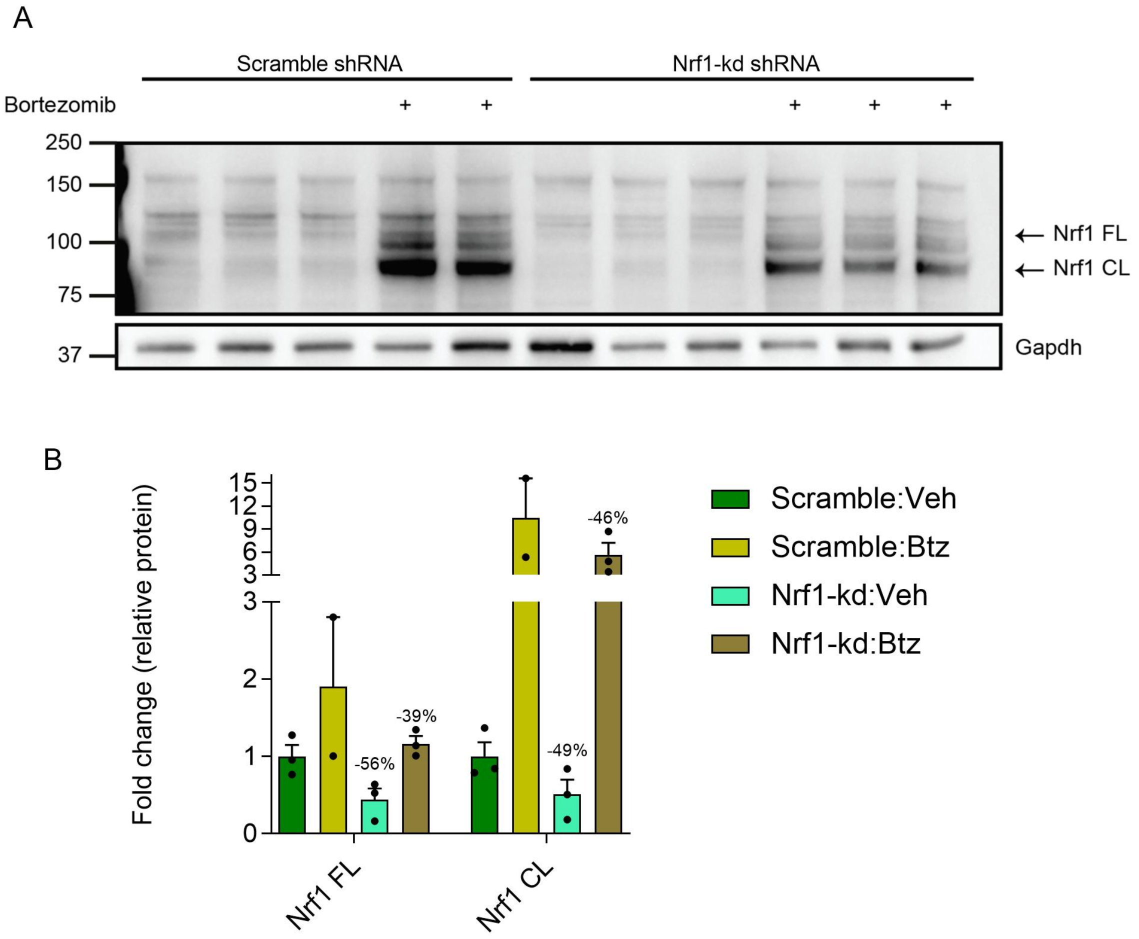


**Figure S5: (A)** Western blots and quantification of cleaved (CL) and full length (FL) Nrf1 in mouse embryonic fibroblasts transfected with plasmids driving the expression of an shRNA directed to Nrf1 (Nrf1-kd) or to a scramble sequence (Scramble) and treated with Bortezomib (BTZ) for 4 hours prior to lysis. Data are presented as mean ± SEM. Percentage change from the respective scramble control is reported for each Nrf1-kd condition. Statistics were not performed due to low sample numbers.

#### SUPPLEMENTARY TABLES

#### Table S1: Unsupervised list of atrophy-related genes

| Gene | Reference | Publication link | Type of muscle atrophy | Category | Direction |
|---|---|---|---|---|---|
| Aco2 | Raffaello et al. (2006) | https://doi.org/10.1152/physiolgenomics.00051.2005 | Denervation | Miscellaneous | Down |
| Actn3 | Raffaello et al. (2006) | https://doi.org/10.1152/physiolgenomics.00051.2005 | Denervation | Miscellaneous | Up |
| Acvr2b | Magnusson et al. (2005) | https://doi.org/10.1111/j.1460-9568.2005.03855.x | Denervation; Negative regulator of muscle growth | Signaling | Down |
| Adam19 | Magnusson et al. (2005) | https://doi.org/10.1111/j.1460-9568.2005.03855.x | Denervation | Signaling | Up |
| Adss | Raffaello et al. (2006) | https://doi.org/10.1152/physiolgenomics.00051.2005 | Denervation | Miscellaneous | Down |
| Ak1 | Raffaello et al. (2006) | https://doi.org/10.1152/physiolgenomics.00051.2005 | Denervation | Miscellaneous | Down |
| Ampd3 | Lecker et al. (2004) | https://doi.org/10.1096/fj.03-0610com | Fasting, Uremia, Tumor, Diabetes | Energy production | Up |
| Apln | Coelho et al. (2019) | https://doi.org/10.1016/j.abb.2019.01.009 | Castration | Miscellaneous | Up |
| Asb15 | Coelho et al. (2019) | https://doi.org/10.1016/j.abb.2019.01.009 | Castration | Miscellaneous | Up |
| Asb2 | Bello et al. (2009) | https://doi.org/10.1038/cdd.2009.27 | no atrophy condition - degradation of filamin B | Protein degradation | Up |
| Atf4 | Lecker et al. (2004) | https://doi.org/10.1096/fj.03-0610com | Fasting, Uremia, Tumor, Diabetes, Den, SC isolation | Transcription | Up |
| Atf6b | Sacheck et al. (2007) | https://doi.org/10.1096/fj.06-6604com | Denervation, Spinal cord isolation | Transcription | Up |
| Atp2a1 | Raffaello et al. (2006) | https://doi.org/10.1152/physiolgenomics.00051.2005 | Denervation | Miscellaneous | Down |
| Atp5a1 | Lecker et al. (2004) | https://doi.org/10.1096/fj.03-0610com | Fasting, Uremia, Tumor, Diabetes, Den, SC isolation | Energy production | Down |
| Atp5b | Raffaello et al. (2006) | https://doi.org/10.1152/physiolgenomics.00051.2005 | Denervation | Miscellaneous | Down |
| Bcat2 | Shimizu et al. (2011) | https://doi.org/10.1016/j.cmet.2011.01.001 | Glucocorticoid induced wasting | Miscellaneous | Up |
| Becn1 | Mammucari et a. (2017) | https://doi.org/10.1016/j.cmet.2007.11.001 | Denervation | Protein degradation | Up |
| Bin1 | Raffaello et al. (2006) | https://doi.org/10.1152/physiolgenomics.00051.2005 | Denervation | Miscellaneous | Up |
| Bnip3 | Lecker et al. (2004) | https://doi.org/10.1096/fj.03-0610com | Fasting, Uremia, Tumor, Diabetes | Protein degradation | Up |
| Cacng1 | Magnusson et al. (2005) | https://doi.org/10.1111/j.1460-9568.2005.03855.x | Denervation | Signaling | Up |
| Cald1 | Raffaello et al. (2006) | https://doi.org/10.1152/physiolgenomics.00051.2005 | Denervation | Miscellaneous | Down |
| Camk2d | Magnusson et al. (2005) | https://doi.org/10.1111/j.1460-9568.2005.03855.x | Denervation | Signaling | Up |
| Cap2 | Sacheck et al. (2007) | https://doi.org/10.1096/fj.06-6604com | Denervation, Spinal cord isolation | Neuromus. Membrane-assoc. | Up |
| Car3 | Raffaello et al. (2006) | https://doi.org/10.1152/physiolgenomics.00051.2005 | Denervation | Miscellaneous | Up |
| Carm1 | Sacheck et al. (2007) | https://doi.org/10.1096/fj.06-6604com | Denervation, Spinal cord isolation | Transcription | Up |
| Cblb | Abe et al. (2013) | http://dx.doi.org/10.1155/2013/907565 | Unloading | Protein degradation | Up |
| Cd68 | Sacheck et al. (2007) | https://doi.org/10.1096/fj.06-6604com | Denervation, Spinal cord isolation | Neuromus. Membrane-assoc. | Up |
| Cd82 | Magnusson et al. (2005) | https://doi.org/10.1111/j.1460-9568.2005.03855.x | Denervation | Signaling | Up |
| Cdkn1a | Magnusson et al. (2005) | https://doi.org/10.1111/j.1460-9568.2005.03855.x | Denervation | Signaling | Up |
| Cdr2 | Magnusson et al. (2005) | https://doi.org/10.1111/j.1460-9568.2005.03855.x | Denervation | Transcription | Up |
| Cebpb | Schakman et al. (2008) | https://doi.org/10.1677/JOE-07-0606 | Dexamethasone | Miscellaneous | Up |
| Cggbp1 | Raffaello et al. (2006) | https://doi.org/10.1152/physiolgenomics.00051.2005 | Denervation | Miscellaneous | Down |
| Chchd10 | Raffaello et al. (2006) | https://doi.org/10.1152/physiolgenomics.00051.2005 | Denervation | Miscellaneous | Down |
| Chrna1 | Sacheck et al. (2007) | https://doi.org/10.1096/fj.06-6604com | Denervation, Spinal cord isolation | Neuromus. Membrane-assoc. | Up |
| Ckmt2 | Lecker et al. (2004) | https://doi.org/10.1096/fj.03-0610com | Fasting, Uremia, Tumor, Diabetes, Den, SC isolation | Energy production | Down |
| Col15a1 | Lecker et al. (2004) | https://doi.org/10.1096/fj.03-0610com | Fasting, Uremia, Tumor, Diabetes | Extracellular matrix | Down |
| Col1a1 | Lecker et al. (2004) | https://doi.org/10.1096/fj.03-0610com | Fasting, Uremia, Tumor, Diabetes, Den, SC isolation | Extracellular matrix | Down |
| Col3a1 | Lecker et al. (2004) | https://doi.org/10.1096/fj.03-0610com | Fasting, Uremia, Tumor, Diabetes | Extracellular matrix | Down |
| Col5a2 | Lecker et al. (2004) | https://doi.org/10.1096/fj.03-0610com | Fasting, Uremia, Tumor, Diabetes, Den, SC isolation | Extracellular matrix | Down |
| Cox7b | Sacheck et al. (2007) | https://doi.org/10.1096/fj.06-6604com | Denervation, Spinal cord isolation | Energy production | Down |
| Cpe | Lecker et al. (2004) | https://doi.org/10.1096/fj.03-0610com | Fasting, Uremia, Tumor, Diabetes, Den, SC isolation | Miscellaneous | Down |
| Cryab | Raffaello et al. (2006) | https://doi.org/10.1152/physiolgenomics.00051.2005 | Denervation | Miscellaneous | Up |
| Cs | Raffaello et al. (2006) | https://doi.org/10.1152/physiolgenomics.00051.2005 | Denervation | Miscellaneous | Down |
| Csnk2a2 | Lecker et al. (2004) | https://doi.org/10.1096/fj.03-0610com | Fasting, Uremia, Tumor, Diabetes, Den, SC isolation | Miscellaneous | Up |
| Csrp3 | Raffaello et al. (2006) | https://doi.org/10.1152/physiolgenomics.00051.2005 | Denervation | Miscellaneous | Up |

| | | | | | |
|---|---|---|---|---|---|
| Ctgf | Magnusson et al. (2005) | https://doi.org/10.1111/j.1460-9568.2005.03855.x | Denervation | Signaling | Up |
| Ctsl | Lecker et al. (2004) | https://doi.org/10.1096/fj.03-0610com | Fasting, Uremia, Tumor, Diabetes | Protein degradation | Up |
| Ctso | Raffaello et al. (2006) | https://doi.org/10.1152/physiolgenomics.00051.2005 | Denervation | Miscellaneous | Down |
| Cyr61 | Magnusson et al. (2005) | https://doi.org/10.1111/j.1460-9568.2005.03855.x | Denervation | Signaling | Up |
| Ddit4 | Shimizu et al. (2011) | https://doi.org/10.1016/j.cmet.2011.01.001 | Glucocorticoid induced wasting | Target of Glucocorticoid receptor | Up |
| Ddx42 | Lecker et al. (2004) | https://doi.org/10.1096/fj.03-0610com | Fasting, Uremia, Tumor, Diabetes | Translation | Up |
| Ddx6 | Sacheck et al. (2007) | https://doi.org/10.1096/fj.06-6604com | Denervation, Spinal cord isolation | Translation | Up |
| Dera | Lecker et al. (2004) | https://doi.org/10.1096/fj.03-0610com | Fasting, Uremia, Tumor, Diabetes | Energy production | Up |
| Dhrs7c | Raffaello et al. (2006) | https://doi.org/10.1152/physiolgenomics.00051.2005 | Denervation | Miscellaneous | Down |
| Dlat | Lecker et al. (2004) | https://doi.org/10.1096/fj.03-0610com | Fasting, Uremia, Tumor, Diabetes, Den, SC isolation | Energy production | Down |
| Dusp5 | Coelho et al. (2019) | https://doi.org/10.1016/j.abb.2019.01.009 | Castration | Miscellaneous | Up |
| Dysf | Sacheck et al. (2007) | https://doi.org/10.1096/fj.06-6604com | Denervation, Spinal cord isolation | Neuromus. Membrane-assoc. | Up |
| Eef1a1 | Raffaello et al. (2006) | https://doi.org/10.1152/physiolgenomics.00051.2005 | Denervation | Miscellaneous | Up |
| Eftud2 | Sacheck et al. (2007) | https://doi.org/10.1096/fj.06-6604com | Denervation, Spinal cord isolation | Transcription | Up |
| Eif4a2 | Lecker et al. (2004) | https://doi.org/10.1096/fj.03-0610com | Fasting, Uremia, Tumor, Diabetes, Den, SC isolation | Translation | Up |
| eIF4E3 | Soares et al. (2014) | https://doi.org/10.1074/jbc.M114.561845 | Fasting, Denervation, Diabetes, Cancer (miRNA analysis) | Miscellaneous | Down |
| Eif4ebp1 | Lecker et al. (2004) | https://doi.org/10.1096/fj.03-0610com | Fasting, Uremia, Tumor, Diabetes, Den, SC isolation | Translation | Up |
| Eif4g2 | Lecker et al. (2004) | https://doi.org/10.1096/fj.03-0610com | Fasting, Uremia, Tumor, Diabetes, Den, SC isolation | Translation | Up |
| Eno3 | Lecker et al. (2004) | https://doi.org/10.1096/fj.03-0610com | Fasting, Uremia, Tumor, Diabetes | Energy production | Down |
| Ep300 | Schakman et al. (2008) | https://doi.org/10.1677/JOE-07-0606 | Dexamethasone | Miscellaneous | Up |
| Ezh1 | Lecker et al. (2004) | https://doi.org/10.1096/fj.03-0610com | Fasting, Uremia, Tumor, Diabetes, Den, SC isolation | Transcription | Up |
| Fam134b | Lecker et al. (2004) | https://doi.org/10.1096/fj.03-0610com | Fasting, Uremia, Tumor, Diabetes | Miscellaneous | Up |
| Fbn1 | Lecker et al. (2004) | https://doi.org/10.1096/fj.03-0610com | Fasting, Uremia, Tumor, Diabetes, Den, SC isolation | Extracellular matrix | Down |
| Fbxo21 | Milan et al. (2015) | https://doi.org/10.1038/ncomms7670 | Starvation, Denervation | Protein degradation | Up |
| Fbxo30 | Sartori et al. (2013) | https://doi.org/10.1038/ng.2772 | Inhibition of BMP sig. (Smad4 KO), Den., Starvation | Protein degradation | Up |
| Fbxo31 | Milan et al. (2015) | https://doi.org/10.1038/ncomms7670 | Starvation, Denervation | Protein degradation | Up |
| Fbxo32 | Lecker et al. (2004) | https://doi.org/10.1096/fj.03-0610com | Fasting, Uremia, Tumor, Diabetes, Den, SC isolation | Protein degradation | Up |
| Fbxo40 | Shi et al. (2011) | https://doi.org/10.1016/j.devcel.2011.09.011 | knockdown induces hypertrophy, degradation of IRS1 | Protein degradation | Up |
| Fbxw7 | Shin et al. (2018) | https://doi.org/10.1007/s11033-018-4185-9 | Dexamethasone | Protein degradation | Up |
| Fhl1 | Raffaello et al. (2006) | https://doi.org/10.1152/physiolgenomics.00051.2005 | Denervation | Miscellaneous | Up |
| Fn1 | Lecker et al. (2004) | https://doi.org/10.1096/fj.03-0610com | Fasting, Uremia, Tumor, Diabetes, Den, SC isolation | Extracellular matrix | Down |
| Foxo1 | Lecker et al. (2004) | https://doi.org/10.1096/fj.03-0610com | Fasting, Uremia, Tumor, Diabetes, Den, SC isolation | Transcription | Up |
| Foxo3 | Ikeda et al. (2016) | https://doi.org/10.1016/j.jtemb.2016.01.011 | Iron-induced | Transcription | Up |
| Frat2 | Magnusson et al. (2005) | https://doi.org/10.1111/j.1460-9568.2005.03855.x | Denervation | Signaling | Down |
| Fst | Schiaffino et al. (2013) | https://doi.org/10.1111/febs.12253 | Inhibits myostatin signaling, induces hypertrophy | Miscellaneous | Down |
| Fth1 | Raffaello et al. (2006) | https://doi.org/10.1152/physiolgenomics.00051.2005 | Denervation | Miscellaneous | Up |
| Fzd9 | Magnusson et al. (2005) | https://doi.org/10.1111/j.1460-9568.2005.03855.x | Denervation | Signaling | Down |
| Gabarapl1 | Lecker et al. (2004) | https://doi.org/10.1096/fj.03-0610com | Fasting, Uremia, Tumor, Diabetes | Protein degradation | Up |
| Gadd45a | Raffaello et al. (2006) | https://doi.org/10.1152/physiolgenomics.00051.2005 | Denervation | Miscellaneous | Up |
| Gclm | Sacheck et al. (2007) | https://doi.org/10.1096/fj.06-6604com | Denervation, Spinal cord isolation | Energy production | Up |
| Glul | Lecker et al. (2004) | https://doi.org/10.1096/fj.03-0610com | Fasting, Uremia, Tumor, Diabetes | Energy production | Up |
| Gpd1 | Lecker et al. (2004) | https://doi.org/10.1096/fj.03-0610com | Fasting, Uremia, Tumor, Diabetes | Energy production | Down |
| Gstm1 | Raffaello et al. (2006) | https://doi.org/10.1152/physiolgenomics.00051.2005 | Denervation | Miscellaneous | Up |
| H3f3a | Raffaello et al. (2006) | https://doi.org/10.1152/physiolgenomics.00051.2005 | Denervation | Miscellaneous | Up |
| Hadh | Sacheck et al. (2007) | https://doi.org/10.1096/fj.06-6604com | Denervation, Spinal cord isolation | Energy production | Down |
| Hectd1 | Lecker et al. (2004) | https://doi.org/10.1096/fj.03-0610com | Fasting, Uremia, Tumor, Diabetes | Miscellaneous | Up |
| Hes6 | Raffaello et al. (2006) | https://doi.org/10.1152/physiolgenomics.00051.2005 | Denervation | Miscellaneous | Down |
| Hmox1 | Kang et al. (2013) | https://doi.org/10.1016/j.febslet.2013.11.009 | Denervation | Oxidative stress | Up |
| Hopx | Coelho et al. (2019) | https://doi.org/10.1016/j.abb.2019.01.009 | Castration | Miscellaneous | Down |
| Hspb1 | Raffaello et al. (2006) | https://doi.org/10.1152/physiolgenomics.00051.2005 | Denervation | Miscellaneous | Up |
| Id2 | Magnusson et al. (2005) | https://doi.org/10.1111/j.1460-9568.2005.03855.x | Denervation | Transcription | Up |
| Idh3b | Sacheck et al. (2007) | https://doi.org/10.1096/fj.06-6604com | Denervation, Spinal cord isolation | Energy production | Down |
| Ifrd1 | Lecker et al. (2004) | https://doi.org/10.1096/fj.03-0610com | Fasting, Uremia, Tumor, Diabetes | Miscellaneous | Up |
| Igf1 | Coelho et al. (2019) | https://doi.org/10.1016/j.abb.2019.01.009 | Castration | Miscellaneous | Down |

| | | | | | |
|---|---|---|---|---|---|
| Igfbp3 | Coelho et al. (2019) | https://doi.org/10.1016/j.abb.2019.01.009 | Castration | Miscellaneous | Up |
| Igfbp5 | Lecker et al. (2004) | https://doi.org/10.1096/fj.03-0610com | Fasting, Uremia, Tumor, Diabetes | Extracellular matrix | Down |
| Impdh2 | Lecker et al. (2004) | https://doi.org/10.1096/fj.03-0610com | Fasting, Uremia, Tumor, Diabetes, Den, SC isolation | Energy production | Up |
| Itch | Milan et al. (2015) | https://doi.org/10.1038/ncomms7670 | Denervation | Protein degradation | Up |
| Itm2a | Lecker et al. (2004) | https://doi.org/10.1096/fj.03-0610com | Fasting, Uremia, Tumor, Diabetes, Den, SC isolation | Miscellaneous | Down |
| Jph2 | Lecker et al. (2004) | https://doi.org/10.1096/fj.03-0610com | Fasting, Uremia, Tumor, Diabetes | Miscellaneous | Down |
| Junb | Lecker et al. (2004) | https://doi.org/10.1096/fj.03-0610com | Fasting, Uremia, Tumor, Diabetes, Den, SC isolation | Transcription | Down |
| Kcna7 | Magnusson et al. (2005) | https://doi.org/10.1111/j.1460-9568.2005.03855.x | Denervation | Signaling | Down |
| Kcnj12 | Magnusson et al. (2005) | https://doi.org/10.1111/j.1460-9568.2005.03855.x | Denervation | Signaling | Down |
| Kcnn3 | Sacheck et al. (2007) | https://doi.org/10.1096/fj.06-6604com | Denervation, Spinal cord isolation | Neuromus. Membrane-assoc. | Up |
| Kctd9 | Lecker et al. (2004) | https://doi.org/10.1096/fj.03-0610com | Fasting, Uremia, Tumor, Diabetes | Energy production | Up |
| Kif21b | Sacheck et al. (2007) | https://doi.org/10.1096/fj.06-6604com | Denervation, Spinal cord isolation | Neuromus. Membrane-assoc. | Down |
| Klf15 | Coelho et al. (2019) | https://doi.org/10.1016/j.abb.2019.01.009 | Castration, Glucocorticoids | Target of Glucocorticoid receptor | Up |
| Klhl38 | Coelho et al. (2019) | https://doi.org/10.1016/j.abb.2019.01.009 | Castration | Miscellaneous | Up |
| Larp7 | Raffaello et al. (2006) | https://doi.org/10.1152/physiolgenomics.00051.2005 | Denervation | Miscellaneous | Down |
| Ldb3 | Lecker et al. (2004) | https://doi.org/10.1096/fj.03-0610com | Fasting, Uremia, Tumor, Diabetes | Miscellaneous | Down |
| Ldha | Lecker et al. (2004) | https://doi.org/10.1096/fj.03-0610com | Fasting, Uremia, Tumor, Diabetes, Den, SC isolation | Energy production | Down |
| Lgals1 | Lecker et al. (2004) | https://doi.org/10.1096/fj.03-0610com | Fasting, Uremia, Tumor, Diabetes, Den, SC isolation | Extracellular matrix | Down |
| Lpin1 | Lecker et al. (2004) | https://doi.org/10.1096/fj.03-0610com | Fasting, Uremia, Tumor, Diabetes | Miscellaneous | Up |
| Lpp | Raffaello et al. (2006) | https://doi.org/10.1152/physiolgenomics.00051.2005 | Denervation | Miscellaneous | Up |
| Luc7l3 | Sacheck et al. (2007) | https://doi.org/10.1096/fj.06-6604com | Denervation, Spinal cord isolation | Miscellaneous | Down |
| Maf | Lecker et al. (2004) | https://doi.org/10.1096/fj.03-0610com | Fasting, Uremia, Tumor, Diabetes | Transcription | Down |
| Maged1 | Magnusson et al. (2005) | https://doi.org/10.1111/j.1460-9568.2005.03855.x | Denervation | Signaling | Up |
| Map1lc3a | Raffaello et al. (2006) | https://doi.org/10.1152/physiolgenomics.00051.2005 | Denervation | Miscellaneous | Up |
| Map1lc3b | Lecker et al. (2004) | https://doi.org/10.1096/fj.03-0610com | Fasting, Uremia, Tumor, Diabetes | Protein degradation | Up |
| Max | Lecker et al. (2004) | https://doi.org/10.1096/fj.03-0610com | Fasting, Uremia, Tumor, Diabetes | Transcription | Up |
| Mb | Raffaello et al. (2006) | https://doi.org/10.1152/physiolgenomics.00051.2005 | Denervation | Miscellaneous | Up |
| Mbnl1 | Raffaello et al. (2006) | https://doi.org/10.1152/physiolgenomics.00051.2005 | Denervation | Miscellaneous | Down |
| Mbtd1 | Raffaello et al. (2006) | https://doi.org/10.1152/physiolgenomics.00051.2005 | Denervation | Miscellaneous | Down |
| Mdh1 | Lecker et al. (2004) | https://doi.org/10.1096/fj.03-0610com | Fasting, Uremia, Tumor, Diabetes, Den, SC isolation | Energy production | Down |
| Mdh2 | Raffaello et al. (2006) | https://doi.org/10.1152/physiolgenomics.00051.2005 | Denervation | Miscellaneous | Down |
| Mdm2 | Ramos et al. (2017) | https://doi.org/10.1111/apha.13003 | T3 (triiodothyronine) stim. (Sarcopenia-Like muscle reg.) | Protein degradation | Up |
| Mkl1 | Coelho et al. (2019) | https://doi.org/10.1016/j.abb.2019.01.009 | Castration | Miscellaneous | Up |
| Mmp14 | Raffaello et al. (2006) | https://doi.org/10.1152/physiolgenomics.00051.2005 | Denervation | Miscellaneous | Up |
| Mstn | Schiaffino et al. (2013) | https://doi.org/10.1111/febs.12253 | Myostatin sig., neg. reg. muscle growth, induces atrophy | Miscellaneous | Up |
| Mt1 | Lecker et al. (2004) | https://doi.org/10.1096/fj.03-0610com | Fasting, Uremia, Tumor, Diabetes | Miscellaneous | Up |
| Mt2 | Magnusson et al. (2005) | https://doi.org/10.1111/j.1460-9568.2005.03855.x | Denervation | Signaling | Up |
| mt-Co1 | Raffaello et al. (2006) | https://doi.org/10.1152/physiolgenomics.00051.2005 | Denervation | Miscellaneous | Down |
| mt-Cytb | Raffaello et al. (2006) | https://doi.org/10.1152/physiolgenomics.00051.2005 | Denervation | Miscellaneous | Down |
| mt-Nd2 | Raffaello et al. (2006) | https://doi.org/10.1152/physiolgenomics.00051.2005 | Denervation | Miscellaneous | Down |
| mt-Nd5 | Raffaello et al. (2006) | https://doi.org/10.1152/physiolgenomics.00051.2005 | Denervation | Miscellaneous | Down |
| Mul1 | Lokireddy et al. (2012) - retracted ("Figure 1D not") | https://doi.org/10.1016/j.cmet.2012.10.005 | Fasting, Denervation | Protein degradation | Up |
| Mybpc1 | Raffaello et al. (2006) | https://doi.org/10.1152/physiolgenomics.00051.2005 | Denervation | Miscellaneous | Down |
| Mybpc2 | Raffaello et al. (2006) | https://doi.org/10.1152/physiolgenomics.00051.2005 | Denervation | Miscellaneous | Down |
| Mybph | Raffaello et al. (2006) | https://doi.org/10.1152/physiolgenomics.00051.2005 | Denervation | Miscellaneous | Up |
| Myc | Magnusson et al. (2005) | https://doi.org/10.1111/j.1460-9568.2005.03855.x | Denervation | Transcription | Up |
| Myh4 | Raffaello et al. (2006) | https://doi.org/10.1152/physiolgenomics.00051.2005 | Denervation | Miscellaneous | Down |
| Myl12a | Raffaello et al. (2006) | https://doi.org/10.1152/physiolgenomics.00051.2005 | Denervation | Miscellaneous | Up |
| Myl2 | Lecker et al. (2004) | https://doi.org/10.1096/fj.03-0610com | Fasting, Uremia, Tumor, Diabetes | Miscellaneous | Down |
| Myog | Sacheck et al. (2007) | https://doi.org/10.1096/fj.06-6604com | Denervation, Spinal cord isolation | Transcription | Up |

| | | | | | |
|---|---|---|---|---|---|
| Myom2 | Raffaello et al. (2006) | https://doi.org/10.1152/physiolgenomics.00051.2005 | Denervation | Miscellaneous | Down |
| Myoz1 | Raffaello et al. (2006) | https://doi.org/10.1152/physiolgenomics.00051.2005 | Denervation | Miscellaneous | Down |
| Ncam1 | Sacheck et al. (2007) | https://doi.org/10.1096/fj.06-6604com | Denervation, Spinal cord isolation | Neuromus. Membrane-assoc. | Up |
| Ncl | Lecker et al. (2004) | https://doi.org/10.1096/fj.03-0610com | Fasting, Uremia, Tumor, Diabetes | Translation | Up |
| Ndufa5 | Sacheck et al. (2007) | https://doi.org/10.1096/fj.06-6604com | Denervation, Spinal cord isolation | Energy production | Down |
| Ndufb5 | Lecker et al. (2004) | https://doi.org/10.1096/fj.03-0610com | Fasting, Uremia, Tumor, Diabetes | Energy production | Down |
| Ndufb8 | Lecker et al. (2004) | https://doi.org/10.1096/fj.03-0610com | Fasting, Uremia, Tumor, Diabetes | Energy production | Down |
| Ndufs1 | Lecker et al. (2004) | https://doi.org/10.1096/fj.03-0610com | Fasting, Uremia, Tumor, Diabetes, Den, SC isolation | Energy production | Down |
| Ndufv2 | Lecker et al. (2004) | https://doi.org/10.1096/fj.03-0610com | Fasting, Uremia, Tumor, Diabetes, Den, SC isolation | Energy production | Down |
| Nedd4 | McClung et al. (2010) | https://doi.org/10.1152/ajpcell.00192.2009 | Oxidative Stress (H2O2) | Oxidative stress | Up |
| Nfe2l2 | Lecker et al. (2004) | https://doi.org/10.1096/fj.03-0610com | Fasting, Uremia, Tumor, Diabetes | Transcription | Up |
| Nptn | Lecker et al. (2004) | https://doi.org/10.1096/fj.03-0610com | Fasting, Uremia, Tumor, Diabetes | Miscellaneous | Up |
| Nr1d1 | Mayeuf-Louchart et al. (2017) | https://doi.org/10.1038/s41598-017-14596-2 | | Miscellaneous | Down |
| Nr4a1 | Magnusson et al. (2005) | https://doi.org/10.1111/j.1460-9568.2005.03855.x | Denervation | Transcription | Down |
| Nrep | Lecker et al. (2004) | https://doi.org/10.1096/fj.03-0610com | Fasting, Uremia, Tumor, Diabetes, Den, SC isolation | Miscellaneous | Down |
| Nsfl1c | Piccirillo et al. (2012) | https://doi.org/10.1038/emboj.2012.178 | Denervation | Protein degradation | Up |
| Oxct1 | Lecker et al. (2004) | https://doi.org/10.1096/fj.03-0610com | Fasting, Uremia, Tumor, Diabetes, Den, SC isolation | Energy production | Down |
| Pdcd10 | Soares et al. (2014) | https://doi.org/10.1074/jbc.M114.561845 | Fasting, Denervation, Diabetes, Cancer (miRNA analysis) | Miscellaneous | Down |
| Pdk4 | Schakman et al. (2008) | https://doi.org/10.1677/JOE-07-0606 | Dexamethasone | Miscellaneous | Up |
| Pfkfb3 | Lecker et al. (2004) | https://doi.org/10.1096/fj.03-0610com | Fasting, Uremia, Tumor, Diabetes | Energy production | Up |
| Pgam2 | Lecker et al. (2004) | https://doi.org/10.1096/fj.03-0610com | Fasting, Uremia, Tumor, Diabetes, Den, SC isolation | Energy production | Down |
| Pgf | Coelho et al. (2019) | https://doi.org/10.1016/j.abb.2019.01.009 | Castration | Miscellaneous | Down |
| Phkg1 | Lecker et al. (2004) | https://doi.org/10.1096/fj.03-0610com | Fasting, Uremia, Tumor, Diabetes, Den, SC isolation | Energy production | Down |
| Pi15 | Coelho et al. (2019) | https://doi.org/10.1016/j.abb.2019.01.009 | Castration | Miscellaneous | Down |
| Pick1 | Magnusson et al. (2005) | https://doi.org/10.1111/j.1460-9568.2005.03855.x | Denervation | Signaling | Down |
| Pik3c3 | Schiaffino et al. (2013) | https://doi.org/10.1111/febs.12253 | Oxidative Stress - nNOS relocalization | Miscellaneous | Up |
| Pik3ip1 | Coelho et al. (2019) | https://doi.org/10.1016/j.abb.2019.01.009 | Castration | Miscellaneous | Up |
| Por | Sacheck et al. (2007) | https://doi.org/10.1096/fj.06-6604com | Denervation, Spinal cord isolation | Miscellaneous | Up |
| Postn | Lecker et al. (2004) | https://doi.org/10.1096/fj.03-0610com | Fasting, Uremia, Tumor, Diabetes, Den, SC isolation | Extracellular matrix | Down |
| Ppargc1a | Wang et al. (2016) | https://doi.org/10.1080/09168451.2016.1254531 | Unloading | Miscellaneous | Down |
| Ppp1r15a | Lecker et al. (2004) | https://doi.org/10.1096/fj.03-0610com | Fasting, Uremia, Tumor, Diabetes | Miscellaneous | Up |
| Ppp5c | Sacheck et al. (2007) | https://doi.org/10.1096/fj.06-6604com | Denervation, Spinal cord isolation | Transcription | Up |
| Prmt1 | Magnusson et al. (2005) | https://doi.org/10.1111/j.1460-9568.2005.03855.x | Denervation | Transcription | Up |
| Prnp | Sacheck et al. (2007) | https://doi.org/10.1096/fj.06-6604com | Denervation, Spinal cord isolation | Miscellaneous | Up |
| Psma1 | Lecker et al. (2004) | https://doi.org/10.1096/fj.03-0610com | Fasting, Uremia, Tumor, Diabetes, Den, SC isolation | Protein degradation | Up |
| Psma5 | Lecker et al. (2004) | https://doi.org/10.1096/fj.03-0610com | Fasting, Uremia, Tumor, Diabetes | Protein degradation | Up |
| Psmb3 | Lecker et al. (2004) | https://doi.org/10.1096/fj.03-0610com | Fasting, Uremia, Tumor, Diabetes, Den, SC isolation | Protein degradation | Up |
| Psmb4 | Lecker et al. (2004) | https://doi.org/10.1096/fj.03-0610com | Fasting, Uremia, Tumor, Diabetes, Den, SC isolation | Protein degradation | Up |
| Psmc4 | Lecker et al. (2004) | https://doi.org/10.1096/fj.03-0610com | Fasting, Uremia, Tumor, Diabetes | Protein degradation | Up |
| Psmd11 | Lecker et al. (2004) | https://doi.org/10.1096/fj.03-0610com | Fasting, Uremia, Tumor, Diabetes, Den, SC isolation | Protein degradation | Up |
| Psmd8 | Lecker et al. (2004) | https://doi.org/10.1096/fj.03-0610com | Fasting, Uremia, Tumor, Diabetes, Den, SC isolation | Protein degradation | Up |
| Psme4 | Lecker et al. (2004) | https://doi.org/10.1096/fj.03-0610com | Fasting, Uremia, Tumor, Diabetes, Den, SC isolation | Protein degradation | Up |
| Ptges3 | Lecker et al. (2004) | https://doi.org/10.1096/fj.03-0610com | Fasting, Uremia, Tumor, Diabetes | Miscellaneous | Up |
| Ptp4a3 | Magnusson et al. (2005) | https://doi.org/10.1111/j.1460-9568.2005.03855.x | Denervation | Signaling | Down |
| Pvalb | Lecker et al. (2004) | https://doi.org/10.1096/fj.03-0610com | Fasting, Uremia, Tumor, Diabetes, Den, SC isolation | Miscellaneous | Down |
| Pygm | Raffaello et al. (2006) | https://doi.org/10.1152/physiolgenomics.00051.2005 | Denervation | Miscellaneous | Down |
| Ramp1 | Magnusson et al. (2005) | https://doi.org/10.1111/j.1460-9568.2005.03855.x | Denervation | Signaling | Down |
| Rb1 | Magnusson et al. (2005) | https://doi.org/10.1111/j.1460-9568.2005.03855.x | Denervation | Transcription | Up |
| Rpl12 | Lecker et al. (2004) | https://doi.org/10.1096/fj.03-0610com | Fasting, Uremia, Tumor, Diabetes, Den, SC isolation | Translation | Up |
| Rpl3 | Raffaello et al. (2006) | https://doi.org/10.1152/physiolgenomics.00051.2005 | Denervation | Miscellaneous | Up |
| Rplp0 | Raffaello et al. (2006) | https://doi.org/10.1152/physiolgenomics.00051.2005 | Denervation | Miscellaneous | Up |

| | | | | | |
|---|---|---|---|---|---|
| Rplp2 | Raffaello et al. (2006) | https://doi.org/10.1152/physiolgenomics.00051.2005 | Denervation | Miscellaneous | Up |
| Rps18 | Raffaello et al. (2006) | https://doi.org/10.1152/physiolgenomics.00051.2005 | Denervation | Miscellaneous | Up |
| Rps20 | Raffaello et al. (2006) | https://doi.org/10.1152/physiolgenomics.00051.2005 | Denervation | Miscellaneous | Up |
| Rps27a | Lecker et al. (2004) | https://doi.org/10.1096/fj.03-0610com | Fasting, Uremia, Tumor, Diabetes, Den, SC isolation | Protein degradation | Up |
| Rrad | Magnusson et al. (2005) | https://doi.org/10.1111/j.1460-9568.2005.03855.x | Denervation | Signaling | Up |
| Rrp9 | Sacheck et al. (2007) | https://doi.org/10.1096/fj.06-6604com | Denervation, Spinal cord isolation | Translation | Up |
| S1pr2 | Pierucci et al. (2018) | https://doi.org/10.1016/j.bbadis.2018.08.040 | Cachexia, Dexamethasone | Miscellaneous | Up |
| Sap30 | Magnusson et al. (2005) | https://doi.org/10.1111/j.1460-9568.2005.03855.x | Denervation | Transcription | Up |
| Sat1 | Lecker et al. (2004) | https://doi.org/10.1096/fj.03-0610com | Fasting, Uremia, Tumor, Diabetes, Den, SC isolation | Energy production | Up |
| Sesn1 | Lecker et al. (2004) | https://doi.org/10.1096/fj.03-0610com | Fasting, Uremia, Tumor, Diabetes | Miscellaneous | Up |
| Sh3glb1 | Lecker et al. (2004) | https://doi.org/10.1096/fj.03-0610com | Fasting, Uremia, Tumor, Diabetes | Miscellaneous | Down |
| Slc25a26 | Lecker et al. (2004) | https://doi.org/10.1096/fj.03-0610com | Fasting, Uremia, Tumor, Diabetes, Den, SC isolation | Energy production | Down |
| Slc25a4 | Raffaello et al. (2006) | https://doi.org/10.1152/physiolgenomics.00051.2005 | Denervation | Miscellaneous | Down |
| Slc35f5 | Lecker et al. (2004) | https://doi.org/10.1096/fj.03-0610com | Fasting, Uremia, Tumor, Diabetes | Energy production | Up |
| Slc3a1 | Sacheck et al. (2007) | https://doi.org/10.1096/fj.06-6604com | Denervation, Spinal cord isolation | Energy production | Up |
| Slc7a8 | Lecker et al. (2004) | https://doi.org/10.1096/fj.03-0610com | Fasting, Uremia, Tumor, Diabetes | Energy production | Up |
| Sln | Sacheck et al. (2007) | https://doi.org/10.1096/fj.06-6604com | Denervation, Spinal cord isolation | Neuromus. Membrane-assoc. | Up |
| Smoc2 | Lecker et al. (2004) | https://doi.org/10.1096/fj.03-0610com | Fasting, Uremia, Tumor, Diabetes | Extracellular matrix | Down |
| Smpx | Raffaello et al. (2006) | https://doi.org/10.1152/physiolgenomics.00051.2005 | Denervation | Miscellaneous | Up |
| Spns2 | Pierucci et al. (2018) | https://doi.org/10.1016/j.bbadis.2018.08.040 | Cachexia, Dexamethasone | Miscellaneous | Up |
| Sqstm1 | Milan et al. (2015) | https://doi.org/10.1038/ncomms7670 | Starvation, Denervation | Protein degradation | Up |
| Srcap | Lecker et al. (2004) | https://doi.org/10.1096/fj.03-0610com | Fasting, Uremia, Tumor, Diabetes | Transcription | Down |
| Srl | Lecker et al. (2004) | https://doi.org/10.1096/fj.03-0610com | Fasting, Uremia, Tumor, Diabetes | Extracellular matrix | Down |
| Tfrc | Lecker et al. (2004) | https://doi.org/10.1096/fj.03-0610com | Fasting, Uremia, Tumor, Diabetes | Miscellaneous | Down |
| Tgif1 | Lecker et al. (2004) | https://doi.org/10.1096/fj.03-0610com | Fasting, Uremia, Tumor, Diabetes, Den, SC isolation | Transcription | Up |
| Timp1 | Magnusson et al. (2005) | https://doi.org/10.1111/j.1460-9568.2005.03855.x | Denervation | Signaling | Up |
| Tnfrsf12a | Hindi et al. (2013) | https://doi.org/10.1096/fj.13-242123 | Denervation (downregulates Pgc1a) | Proinflammtory cytokine | Up |
| Tnfsf12 | Hindi et al. (2013) | https://doi.org/10.1096/fj.13-242123 | Denervation (downregulates Pgc1a) | Proinflammtory cytokine | Up |
| Tpi1 | Lecker et al. (2004) | https://doi.org/10.1096/fj.03-0610com | Fasting, Uremia, Tumor, Diabetes, Den, SC isolation | Energy production | Down |
| Traf6 | Paul et al. (2010) | https://doi.org/10.1083/jcb.201006098 | Denervation, Diabetes, Cancer | Protein degradation | Up |
| Trib3 | Choi et al. (2019) | https://doi.org/10.1096/fj.201802145RR | Fasting/Starvation | Stress response | Up |
| Trim32 | Kudryashova et al. (2005) | https://doi.org/10.1016/j.jmb.2005.09.068 | Unloading | Protein degradation | Up |
| Trim54 | Centner et al. (2001) | https://doi.org/10.1006/jmbi.2001.4448 | Immobilization, Denervation | Protein degradation | Up |
| Trim55 | Centner et al. (2001) | https://doi.org/10.1006/jmbi.2001.4448 | Immobilization, Denervation | Protein degradation | Up |
| Trim63 | Bodine et al. (2001) | https://doi.org/10.1126/science.1065874 | Immobilization, Denervation | Protein degradation | Up |
| Ttn | Raffaello et al. (2006) | https://doi.org/10.1152/physiolgenomics.00051.2005 | Denervation | Miscellaneous | Down |
| Txnl1 | Lecker et al. (2004) | https://doi.org/10.1096/fj.03-0610com | Fasting, Uremia, Tumor, Diabetes, Den, SC isolation | Miscellaneous | Up |
| Uba52 | Lecker et al. (2004) | https://doi.org/10.1096/fj.03-0610com | Fasting, Uremia, Tumor, Diabetes, Den, SC isolation | Protein degradation | Up |
| Ubb | Lecker et al. (2004) | https://doi.org/10.1096/fj.03-0610com | Fasting, Uremia, Tumor, Diabetes, Den, SC isolation | Protein degradation | Up |
| Ubc | Lecker et al. (2004) | https://doi.org/10.1096/fj.03-0610com | Fasting, Uremia, Tumor, Diabetes, Den, SC isolation | Protein degradation | Up |
| Ube2j1 | Lecker et al. (2004) | https://doi.org/10.1096/fj.03-0610com | Fasting, Uremia, Tumor, Diabetes | Protein degradation | Up |
| Ube3b | Kerksick et al. (2013) | https://doi.org/10.1016/j.fct.2013.01.026 | Exercise - Eccentric muscle contractions | Protein degradation | Up |
| Ube4b | Lecker et al. (2004) | https://doi.org/10.1096/fj.03-0610com | Fasting, Uremia, Tumor, Diabetes | Protein degradation | Up |
| Uqcrc2 | Sacheck et al. (2007) | https://doi.org/10.1096/fj.06-6604com | Denervation, Spinal cord isolation | Energy production | Down |
| Usp14 | Lecker et al. (2004) | https://doi.org/10.1096/fj.03-0610com | Fasting, Uremia, Tumor, Diabetes, Den, SC isolation | Protein degradation | Up |
| Vcp | Piccirillo et al. (2012) | https://doi.org/10.1038/emboj.2012.178 | Denervation, Starvation | Protein degradation | Up |
| Vdac1 | Raffaello et al. (2006) | https://doi.org/10.1152/physiolgenomics.00051.2005 | Denervation | Miscellaneous | Down |
| Vegfa | Magnusson et al. (2005) | https://doi.org/10.1111/j.1460-9568.2005.03855.x | Denervation | Signaling | Down |
| Zbtb16 | Magnusson et al. (2005) | https://doi.org/10.1111/j.1460-9568.2005.03855.x | Denervation | Transcription | Down |

**Table S2: Primers used for quantitative PCR analysis**

| Gene | Forward primer | Reverse primer |
|---|---|---|
| *Actb* | CAGCTTCTTTGCAGCTCCTT | GCAGCGATATCGTCATCCA |
| *Bnip3* | TTCCACTAGCACCTTCTGATGA | GAACACCGCATTTACAGAACAA |
| *Tubb* | GAAGCTGACCACACCCACCT | TGAAGAAGTGCAGGCGTGGG |
| *Ctsd* | CGTCTTGCTGCTCATTCTCGGCC | GCCGCCCACCTCCGTCATAGT |
| *Ctsl* | GTGGACTGTTCTCACGCTCA | TCCGTCCTTCGCTTCATAGG |
| *Des* | GAGGTTGTCAGCGAGGCTAC | CTTCAGGAGGCAGTGAGGAC |
| *Fbxo30* | TCGTGGAATGGTAATCTTGC | CCTCCCGTTTCTCTATCACG |
| *Fbxo31* | GTATGGCGTTTGTGAGAACC | AGCCCCAAAATGTGTCTGTA |
| *Fbxo32* | CTCTGTACCATGCCGTTCCT | GGCTGCTGAACAGATTCTCC |
| *Gabarapl1* | CATCGTGGAGAAGGCTCCTA | ATACAGCTGGCCCATGGTAG |
| *Gadd45a* | CCGAAAGGATGGACACGGTG | TTATCGGGGTCTACGTTGAGC |
| *Map1lc3b* | CACTGCTCTGTCTTGTGTAGGTTG | TCGTTGTGCCTTTATTAGTGCATC |
| *Mdm2* | GAGGATGATGAGGTCTATCG | GGAGGATTCATTTCATTGCAC |
| *Trim63* | ACCTGCTGGTGGAAAACA | AGGAGCAAGTAGGCACCTCA |
| *Nfe2l1* | CCCTACTCACCCAGTCAGTATG | CATCGTGCGAGGAATGAGGA |
| *Nfe2l2* | GCCCACATTCCCAAACAAGAT | CCAGAGAGCTATTGAGGGACTG |
| *Nqo1* | TTCTCTGGCCGATTCAGAG | GGCTGCTTGGAGCAAAATAG |
| *Ogt* | TTCGGGAATCACCCTACTTCA | TACCATCATCCGGGCTCAA |
| *Ppp1r15a* | GACCCCTCCAACTCTCCTTC | CTTCCTCAGCCTCAGCATTC |
| *Psma1* | CCTCAGGGCAGGATTCATCAA | GAGCGGCAAGCTCTGACTG |
| *Psma5* | AGCAATTGGCTCTGCTTCAG | GCATTCAGCTTCTCCTCCAT |
| *Psmb5* | AGGAGCCGCGAATCGAAATG | CCAGAAGGTACGGGTTGATCTC |
| *Psmb6* | CTGGGAAAACCGGGAAGTCTC | GAGTCCGCTCCTAGAACCAC |
| *Psmb7* | GTGTCGGTGTTTCAGCCAC | TCCGCTTCCAAGACAGCATT |
| *Psmc1* | AAGGGGGTCATTCTCTACGG | AAGCTCTGAGCCAACCACTC |
| *Psmc4* | TGGTCATCGGTCAGTTCTTG | CGGTCGATGGTACTCAGGAT |
| *Psmd4* | TCTCCTATTCTGGCTGGTGAA | CATGCTGCTTAGGTCTGGAAG |
| *Psmd8* | GCCTCAATCTCCTCTTCCTGCTATC | GTCTGTCATCTTTTTGGGTGTGC |
| *Psmd11* | AGGCAGACAGAAGCATTGAAA | GGTCCAAAATCCCATGAAACT |
| *Psme4* | AGCGTCAACAAGATAAGAATGCT | GCCCGATTCCTATATGCTCAAA |
| *Sesn1* | CCATAGGCCTTGGCTGATTA | AGCTGTGCTCCTCTGCTTTC |
| *Sqstm1* | AGGGAACACAGCAAGCTCAT | GACTCAGCTGTAGGGCAAGG |
| *Traf6* | GCAGTGAAAGATGACAGCGTGA | TCCCGTAAAGCCATCAAGCA |
| *Ubb* | GCTTACCATGCAACAAAACCT | CCAGTGGGCAGTGATGG |
| *Ube4b* | TGTCATCTTCCTTTCTTCTCTCT | TGGATTTTCATCTCGTGTCTG |
| *Vcp* | GGTTGGGGTTAGAGCAGCTT | GCGACTAATCAAACGACGGC |

**Table S3 Primary antibodies used for immunoblotting**

| Protein | Catalog number | Company | Dilution |
|---|---|---|---|
| 4E-BP1 | #9452 | Cell Signaling Technology | 1:1000 |
| p-4E-BP1 S65 | #9451 | Cell Signaling Technology | 1:1000 |
| a-actinin | A7732 | Sigma | 1:5000 |
| Akt | #9272 | Cell Signaling Technology | 1:1000 |
| p-Akt S473 | #9271 | Cell Signaling Technology | 1:1000 |
| Bnip3 | 3769 | Cell Signaling Technology | 1:1000 |
| GAPDH | #2118 | Cell Signaling Technology | 1:5000 |
| Mono + Polyubiquitin | BML-PW8810 | Enzo | 1:500 |
| Nfe2l1 (Nrf1) | #8052 | Cell Signaling Technology | 1:1000 |
| Ogt | #24083 | Cell Signaling Technology | 1:1000 |
| p62 | GP62-C | Progen | 1:1000 |
| PRAS40 | #2610 | Cell Signaling Technology | 1:1000 |
| p- PRAS40 T246 | #2997 | Cell Signaling Technology | 1:1000 |
| PSMA* | BML-PW8195 | Enzo | 1:1000 |
| PSMB5 | ab3330 | Abcam | 1:1000 |
| PSMB6 | #13267 | Cell Signaling Technology | 1:1000 |
| PSMB7 | #13207 | Cell Signaling Technology | 1:1000 |
| PSMB8 | BML-PW8845 | Enzo | 1:1000 |
| PSCM1 | ab140450 | Abcam | 1:2000 |
| PSMC5 | ab140450 | Abcam | 1:2000 |
| PSME4 | 18799-1-AP | Proteintech | 1:500 |
| Puromycin | MABE343 | Millipore | 1:5000 |
| S6 | #2217 | Cell Signaling Technology | 1:1000 |
| p-S6 S235/S236 | #2211 | Cell Signaling Technology | 1:1000 |
| p-S6 S240/S244 | #5364 | Cell Signaling Technology | 1:1000 |
| SREBF1 | Sc-8984 | Santa Cruz | 1:1000 |
| TSC1 | A300-316A | Bethyl | 1:5000 |
| Vcp (p97) | #2648 | Cell Signaling Technology | 1:1000 |
